# Endothelial cell flow-mediated quiescence is temporally regulated and utilizes the cell cycle inhibitor p27

**DOI:** 10.1101/2023.06.09.544403

**Authors:** Natalie T Tanke, Ziqing Liu, Michaelanthony T Gore, Pauline Bougaran, Mary B Linares, Allison Marvin, Arya Sharma, Morgan Oatley, Tianji Yu, Kaitlyn Quigley, Sarah Vest, Jeanette Gowen Cook, Victoria L Bautch

**Author notes:** Corresponding author: Victoria L Bautch, PhD, Professor of Biology, Department of Biology, CB#3280 University of North Carolina at Chapel Hill Chapel Hill, NC 27599 USA.

## Abstract

**Background:** Endothelial cells regulate their cell cycle as blood vessels remodel and transition to quiescence downstream of blood flow-induced mechanotransduction. Laminar blood flow leads to quiescence, but how flow-mediated quiescence is established and maintained is poorly understood.

**Methods:** Primary human endothelial cells were exposed to laminar flow regimens and gene expression manipulations, and quiescence depth was analyzed via time to cell cycle re-entry after flow cessation. Mouse and zebrafish endothelial expression patterns were examined via scRNA seq analysis, and mutant or morphant fish lacking p27 were analyzed for endothelial cell cycle regulation and *in vivo* cellular behaviors.

**Results:** Arterial flow-exposed endothelial cells had a distinct transcriptome, and they first entered a deep quiescence, then transitioned to shallow quiescence under homeostatic maintenance conditions. In contrast, venous-flow exposed endothelial cells entered deep quiescence early that did not change with homeostasis. The cell cycle inhibitor p27 (*CDKN1B)* was required to establish endothelial flow-mediated quiescence, and expression levels positively correlated with quiescence depth. p27 loss *in vivo* led to endothelial cell cycle upregulation and ectopic sprouting, consistent with loss of quiescence. *HES1* and *ID3*, transcriptional repressors of p27 upregulated by arterial flow, were required for quiescence depth changes and the reduced p27 levels associated with shallow quiescence.

**Conclusions:** Endothelial cell flow-mediated quiescence has unique properties and temporal regulation of quiescence depth that depends on the flow stimulus. These findings are consistent with a model whereby flow-mediated endothelial cell quiescence depth is temporally regulated downstream of p27 transcriptional regulation by HES1 and ID3. The findings are important in understanding endothelial cell quiescence mis-regulation that leads to vascular dysfunction and disease.

**GRAPHICAL ABSTRACT:** 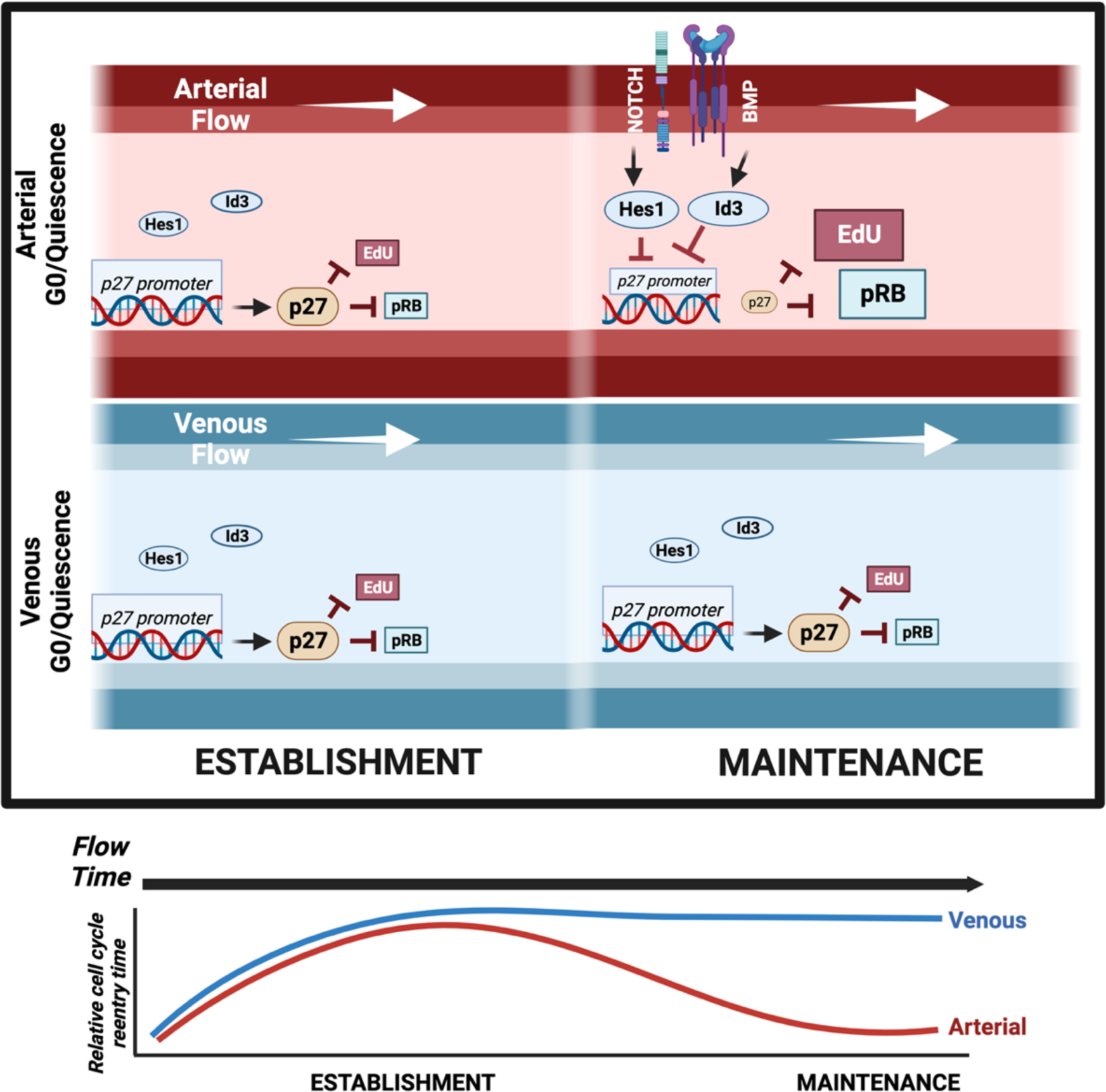

**HIGHLIGHTS:**

- Different quiescence stimuli lead to distinct transcriptional and functional quiescence profiles in endothelial cells
- p27 is required for endothelial cell quiescence and depth is temporally regulated in a flow stimulus-dependent manner that correlates with p27 levels and flow-regulated repressors *HES1* and *ID3*
- p27 is expressed in endothelial cells according to flow magnitude *in vivo* and is functionally required for cell cycle regulation and sprouting *in vivo*

## INTRODUCTION

Developmental blood vessel network expansion via sprouting angiogenesis leads to remodeling and homeostasis; this transition depends on endothelial cell responses to incoming signals, including laminar shear stress provided by blood flow. Laminar flow leads to endothelial cell cytoskeletal realignment and dramatically reduces proliferation^1–6^. Transcriptional profiles change with flow exposure^7,8^, and signaling pathways such as Notch and BMP are upregulated to promote alignment and vascular homeostasis^9,10^. Vascular homeostasis significantly represses endothelial cell proliferation and sets up a quiescent (G_0_) state that can be released by angiogenic growth factors such as VEGF-A^7,11,12^; ^13–15^. How flow-mediated vascular quiescence is established and maintained is poorly understood, yet it is critical to blood vessel function.

Quiescence is influenced by cell type in non-endothelial cells^16,17^, and quiescence stimuli such as growth factor deprivation or contact inhibition also influence quiescence parameters^18^. Different quiescence stimuli lead to transcriptional profiles that are partially overlapping but have strong unique signatures^16,18^; however, how these differences affect quiescence properties is poorly understood, and it is not known how endothelial cells differentially respond to quiescence cues.

The cell cycle is regulated by cyclins and cyclin-dependent kinases (CDKs) that promote and CDK inhibitors that block cell cycle progression^19^. In cycling cells, proteins are post-translationally modified and/or carry destabilization sequences that lead to rapid gain and loss of function required for passage through the cell cycle^20,21^. In contrast, quiescence is characterized by transcriptional repression of cell cycle activators and often by upregulation of cell cycle inhibitors^18,19^. Thus, cell cycle reentry is delayed upon removal of the stimulus, and reentry time is used as a proxy for quiescence depth^22,23^. Quiescence depth is variable; for example, quiescent fibroblasts have delayed cell cycle reentry that positively correlates with temporal exposure to the quiescence stimulus^22,23^. Non-homogeneous quiescence depth is associated with muscle stem cells that respond to injury-induced circulating cues by entering a shallow quiescence called “G_0_ alert” that allows for prompt proliferation in response to a subsequent injury^24^. Moreover, spontaneous quiescence was recently described, whereby epithelial cells enter quiescence absent obvious environmental or pharmacological cues, suggesting that quiescence entry has a stochastic component and/or complex inputs^16,25^. Thus, how quiescence is established, maintained, and exited is complex.

We examined properties of flow-induced endothelial cell quiescence and found that arterial laminar flow-induced endothelial cell quiescence had a distinct transcriptional profile and functionally exhibited dynamic temporal regulation of quiescence depth, while venous flow did not exhibit the same temporal regulation of quiescence depth. Temporal changes in endothelial cell quiescence depth positively correlated with expression of the cell cycle inhibitor p27, and p27 repression and temporal regulation of quiescence depth under arterial flow required *HES1* and *ID3*, targets of the Notch and BMP signaling pathways. A requirement for p27 for endothelial quiescence *in vivo* and correlation of p27 expression with flow magnitude was revealed. Thus, flow-mediated endothelial cell quiescence is novel and complex, and regulation of quiescence depth correlates with flow stimulus and likely influences physiological and pathological vascular quiescence responses.

## MATERIALS AND METHODS

### Data availability

Data associated with this study are available from the corresponding author upon reasonable request. **Key Resources Table** is in Supplementary File.

### Cell culture

HUVEC and HAEC were cultured according to the manufacturer’s recommendations in EBM2 (Lonza#CC-3162) with growth factors (Lonza#CC-3162, EGM2) at 37°C, 5% CO_2_ and used between passages 2-4 (**Key Resources Table, cell culture)**. Critical experiments were replicated with multiple lots of HUVEC and with HAEC.

### Microscopy

Imaging was performed using confocal microscopy (Olympus Fluoview FV3000, IX83) and a UPlanSApo 40x silicone-immersion objective (NA 1.25), UPlanSApo 60x oil-immersion objective (NA 1.40), or UPlanSApo 100x oil-immersion objective (NA 1.40). Images were acquired with Fluoview FV31S-SW software and imaging analysis was completed in FIJI^26^. For each replicate within an experiment, images were acquired at the same settings.

### Endothelial Cell Transfection

HUVEC were grown to sub-confluency and treated with siRNAs **(Key Resources Table, siRNA information)** diluted in OptiMem (Gibco, #11058021) and Lipofectamine 3000 (ThermoFisher, #L3000015) according to manufacturer’s protocol (https://www.thermofisher.com/us/en/home/references/protocols/cell-culture/transfection-protocol/lipofectamine-2000.html). Briefly, siRNA at 0.48 µM in Opti-MEM (31985-070, Gibco) and a 1:20 dilution of Lipofectamine in Opti-MEM were incubated separately at RT for 5 min, then combined and incubated at RT for 15 min. HUVEC were transfected at 80% confluency with siRNA at 37°C for 24h, then in 10 mL of fresh EGM-2. All experiments were initiated 48h following siRNA exposure.

### Immunofluorescence and EdU labeling

Endothelial cells were fixed in 4% PFA (15713 (100504-940), VWR) at 37°C for 10 min, permeabilized in 0.1% Triton (T8787-100ML, Sigma) in PBS for 10 min at RT, then blocked for 1h at RT in 5% NBCS (Gibco, #16010-159), antibiotic-antimycotic (Gibco, #15240062), 0.1% sodium azide (Sigma s2002-100G). Following 3X PBS washes, cells were incubated in primary antibody/blocking solution and incubated for 24h at 4°C (**Key Resources Table, antibodies**), washed 3X with PBS and incubated in secondary antibody/blocking solution for 3h at RT in the dark (**Key Resources Table, antibodies**). Slides were mounted with coverslips using Prolong Diamond Antifade mounting media (P36961, Life Technology), sealed with nail polish, and stored at 4°C. Glass-bottom Ibidi slides or well dishes were stored in 1X PBS. For all nuclear stains, The DAPI channel was used as a nuclear mask, and nuclear fluorescence intensity was measured for each cell per image field. Positive cells were defined as those above the sensitivity threshold.

EdU labeling was performed according to the Click-IT EdU 488, 594, or 647 protocol (Invitrogen, C10337). Cells were incubated in EdU for 30 min (quiescence depth) or 1h at 37°C in 5% CO_2_ and fixed with 4% PFA for 10 min at RT.

### Quiescence Depth

#### Flow

HUVEC were cultured in EBM2 supplemented with 2% FBS, 1% antibiotic-antimycotic (Gibco, #15240062), and 1% Nystatin (Sigma, #N1638-20ML) and exposed to laminar flow for indicated times and shear levels using an Ibidi pump system (CC-1090, Ibidi). Static (non-flow) control slides were seeded at lower densities and incubated for 48h prior to fixation with 4% PFA. For western blot experiments, an orbital shaker (Hoefer Red Rotor Mixer Platform Shaker PR70-115V) was used as previously described^27^. Flow-mediated quiescence depth experiments were performed after 16h or 72h of laminar flow exposure by incubation for 30 min with EdU, fixation at indicated times post-flow in 4% PFA for 10 min at RT, then visualized for EdU and/or stained for antibodies as described **(Supp. Tables S2, S3)**.

#### Contact

HUVEC were seeded in a 6-well plate at 1.2 X 10^6^ cells/well (high density) or 0.2 X 10^6^ cells/well (low density) and incubated for 24h. A p1000 pipette tip was used to scrape the cell monolayer and create a gap, as previously described^28^, cells were incubated with EdU for 30 min or fixed at 0h, 2h, 5h, or 8h post-scratching in 4% PFA for 10 min at RT, then visualized for EdU and/or stained for antibodies as described **(Supp. Table S2, S3)**. Cells within 1000μm of the scratch edge were imaged **(Supp. Fig. 2a)**.

### Cell Axis Ratio

Cell shape and alignment were measured as described previously^7^. Cells were stained for VE-cadherin upon fixation, and measured at the longest axis of cell divided by shortest axis to calculate the cell axis ratio, with the longest cell axis being the direction of flow.

### Western Blot Analysis

Western blot analysis was performed according to^29^ with modifications. Briefly, cells were scraped into PBS, centrifuged at 13000 rpm (4°C, 20 min), resuspended in RIPA buffer with protease/phosphatase inhibitor (5872S, Cell Signaling), then added to sample loading buffer with dithiothreitol (R0861, Thermo Fisher) and boiled for 10 min. Samples (10ug) were separated on 10% SDS-PAGE gel (161-0183, BioRad), then transferred to a membrane, incubated in primary antibodies (overnight, 4°C) (**Key Resources Table, antibodies**), washed 3X in PBST and stained with secondary antibody in One-Block (RT, 1h) (**Key Resources Table, antibodies**). Immobilon Forte HRP Substrate (WBLUF0100, 769 Millipore Sigma) was added for 30 sec, and blots were exposed for 2 sec ChemiDoc XRS with Chemi High Resolution setting.

### RNA

#### RT-qPCR

Scraped cell pellets were resuspended in TRIzol (15596018, Invitrogen). cDNA was generated from 1ug RNA using iScript reverse transcription kit (Bio-Rad, #1708891) and diluted 1:3 in water. qRT–PCR was performed using iTaq Universal SYBR Green SuperMix (Bio-Rad, #1725121). SYBR Green real-time PCR was performed in triplicate on the Applied Biosystems QuantStudio 6 Flex Real-Time PCR System (**Key Resources Table, qPCR primers**). For quantification, relative expression of each gene to β-actin in each sample was calculated by 2^ (CT of gene−CT of β-actin). Statistical significance was determined by unpaired Student’s T-test.

#### Bulk RNA seq

3 Ibidi slides/condition were pooled, and stranded libraries were prepared using KAPA mRNA HyperPrep Kit (7961901001, Roche) and sequenced using NovaSeq S1 at the UNC Sequencing Core. Data was analyzed as described previously ^30^. Briefly, 2–3×10^7^ 50 bp paired-end reads per sample were obtained and mapped to the human genome GRCh38 with STAR using default settings^31^. Quality of sequencing reads was confirmed with FastQC before mapping. Mapping rate was over 87% for all samples **(Supp. Table S2)**, and gene (GEO GSE213323) expression was determined with Htseq-count using the union mode^32^ . Genes with low expression were filtered out (total raw counts in all samples <10), differential expression analysis was performed with DESeq2 ^33^ using default settings in R, and lists of differentially expressed genes were obtained (p adjusted <0.1). Gene ontology analysis was performed using enrichGO function in the R package clusterProfiler. All gene ontology terms shown in this study have a corrected P value <0.1.

Mouse scRNAseq: A mouse scRNAseq dataset (Liu et al, in preparation) (GEO GSE216594) previously generated using enriched endothelial cells from the mouse ear at P8 (postnatal day 8) was analyzed for *Cdkn1b* levels in endothelial cells.

### Quiescence Score

The endothelial quiescence score was calculated using the formula:QS = 1/n∑_*i* = 1_^*n*^ *QSUi*-1/m1/n∑_*j* = 1_^*m*^ *QSDj* (QS = Quiescence Score, QSU = QS of upregulated genes under quiescence (total n genes), QSD = QS of downregulated genes under quiescence (total m genes)). Selected genes were upregulated or downregulated relative to static samples. All genes were normalized for sequencing depth and scaled for expression as previously described^8^. To generate the gene list used for endothelial quiescence score we utilized two datasets **(Supp. Table S1)**: S/G_2_/M phase associated genes from the CellCycleScoring function of the R package Seurat that were down-regulated with flow in scRNAseq HUVEC (accession code GSE151867)^8^, and genes that were upregulated with different quiescence stimuli in non-endothelial cells^18^ and upregulated with flow in scRNAseq data. The epithelial quiescence score was developed by the Barr lab as previously described, and utilized an independent gene list generated using non-transformed human epithelial cell data^16,34^.

### Zebrafish

Zebrafish (*Danio rerio*) were housed in an IACUC approved facility^30^. *Tg(fli:LifeAct-GFP*) was a gift from Dr. Wiebke Herzog **(Key Resources Table, animals)**. The *cdkn1bb* CRISPR line was designed as previously described **(Key Resources Table**, **genetically modified animals)**^35^. One-cell stage zebrafish embryos were injected with 1nl injection solution into the cytoplasm. Fin clips were used to identify possible founders with gene-flanking primers PCR (F-CTCAATAACTGCTGCGAGTG, R-GATGAAGGGGGGAAAGAGG, R-GCCATCGAGTCAAACCAG).

Morphant fish were obtained by randomly injecting 2.5–5 ng of non-targeting (NT) (5’-CCTCTTACCTCAGTTACAATTTATA-3’, GeneTools, LLC) or *cdkn1bb* (5’-ACGGTCAAAATTCAAAGCACATACC 3’, GeneTools, LLC) MO into *Tg(fli:LifeAct-GFP*) embryos at the one-cell stage. Fish were grown in E3 medium at 28.5°C to 36hpf.

Embryos were prepared for imaging as previously described^30^. Briefly, dechorionated embryos were fixed overnight in ice-cold 4% PFA at 4 °C, rinsed 2X in PBS, mounted using Prolong Diamond Antifade mounting medium (P36961, Life Technology), and coverslip sealed with petroleum jelly.

### Zebrafish FACs Sorting

FACs sorting was as previously described^36^. Briefly, embryos were euthanized in 1X Tricaine, dissociated with 100mg/mL collagenase in 1X PBS and 0.25% trypsin and harsh mechanical pipetting, and incubated (30°C, 10 min). The dissociation reaction was neutralized with DMEM + 10% FBS and cells were centrifuged (5000 rcf, 5 min). Pellets were resuspended in 500μL DMEM/FBS and filtered using a 30μm cell strainer. Samples were analyzed on the AttuneX (UNC Flow Cytometry Core), and GFP+ cells from *fli:LifeAct-GFP* embryos were collected into TRIzol.

### Statistics

Unpaired Students T-Test was used to determine statistical significance with two experimental groups, and One-Way ANOVA with Tukey correction was used for experimental groups ≥3. Ξ^2^ test was performed to compare distribution across 2 groups. All statistical tests and graphs were made using the Prism 9.4.1 software (GraphPad Software) and are described in relevant Figure Legends.

## RESULTS

### Laminar flow-exposed endothelial cells have a distinct transcriptional profile

Endothelial cells exposed to laminar shear stress exhibit alignment to the flow vector and reduced proliferation, hallmarks of vascular homeostasis or quiescence^1,2,5^. To determine whether endothelial cells exposed to laminar flow were transcriptionally quiescent, we utilized a published quiescence score formula generated in non-endothelial epithelial cells^34^ and applied it to a scRNAseq dataset we previously generated in primary endothelial cells (HUVEC, human umbilical vein endothelial cells) exposed to laminar flow^8^. This analysis revealed that endothelial cells exposed to homeostatic laminar flow (here called Flow-Maintenance (Flow-M) (15d/cm^2^, 72h)) had a significantly higher score than non-flowed control endothelial cells **(Fig. 1A)**. Previous transcriptional characterization revealed 9 clusters primarily separated by flow condition, and the cluster containing the majority of cells exposed to flow (Flow Cluster 1) had a significantly higher epithelial quiescence score compared to the equivalent control cluster (Static Cluster 1), indicating that this quiescence score positively correlates with flow-exposed endothelial cells predicted to be quiescent. Quiescence score comparisons among minor flow clusters showed heterogeneity relative to Flow Cluster 1, and a similar comparison of static clusters also revealed heterogeneity relative to Static Cluster 1, indicating that distinct clusters exhibit quiescence heterogeneity. As predicted, Flow Cluster 2, Static Cluster 2, and Mix Cluster (endothelial cells from flow and static conditions), previously defined as proliferative populations^8^, had a significantly reduced quiescence score **(Fig. 1A).**

**Figure 1.**
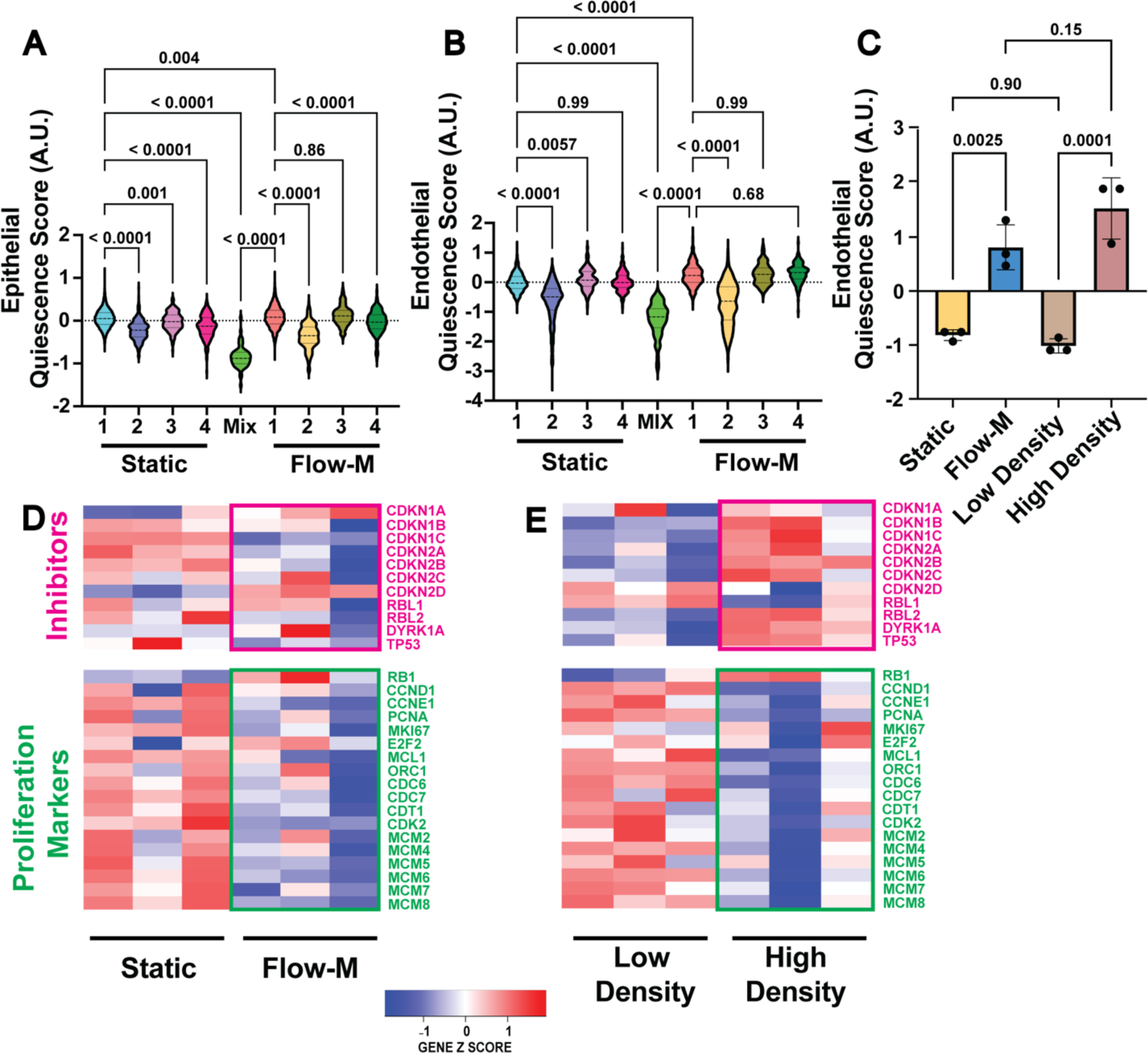
Laminar flow-mediated quiescence is transcriptionally distinct. **(A)** Quantification of epithelial quiescence score in HUVEC scRNA dataset by cluster (previously defined^8^). Flow-M, flow maintenance (laminar flow (15d/cm^2^/72h)). Static cells were visually checked for subconfluence and collected 48h post-seeding. **(B)** Quantification of endothelial quiescence score in HUVEC scRNA dataset by cluster. **(C)** Endothelial quiescence score on bulk RNA seq data of HUVEC exposed to indicated stimuli, n= 3 replicates. **(D-E)** Heatmaps showing relative expression of cell cycle proliferation markers (green font) and inhibitors (red font) plotted using bulk RNAseq data of HUVEC under indicated conditions, n=3 replicates. Statistics, one-way ANOVA with Tukey’s multiple comparison test.

We next used the scRNA seq dataset to generate an independent endothelial quiescence score algorithm. Since a hallmark of cellular quiescence is downregulation of proliferation markers and upregulation of cell cycle inhibitors^18^, this score was primarily generated using expression levels of cell cycle genes that were down- or upregulated in endothelial cells exposed to homeostatic laminar flow (Flow-M) **(Supp. Table 1)**. The endothelial quiescence score was then applied to the dataset to verify that the score predicted the relationships. Consistent with the epithelial score, Flow Cluster 1 had a significantly higher quiescence score compared to Static Cluster 1, and the proliferation-associated clusters associated had significantly lower scores **(Fig. 1B).** These findings indicate that endothelial cells exposed to homeostatic laminar flow (Flow-M) are in a transcriptionally quiescent cell cycle state.

Non-endothelial cells exposed to different quiescence stimuli have unique transcriptional profiles^16,18^, so we hypothesized that the mode of quiescence induction affects the quiescence score of endothelial cells. Application of the endothelial quiescence score algorithm to HUVEC bulk RNAseq data showed significant increases under flow and high density compared to static and low-density conditions, respectively **(Fig. 1C)**, and these relationships held when the epithelial quiescence score algorithm was applied **(Supp. Fig. 1A)**. Venn diagrams revealed that homeostatic laminar flow (Flow-M) and contact inhibition have both distinct and shared transcriptional changes in endothelial cells, with approximately 25% of regulated genes shared for up- and down-regulated categories (upregulated: 21.7% (494/2272) flow and 24.3% (494/2030) density shared genes; down-regulated 25.1% (573/2282) flow and 23.7% (573/2415) density shared genes) **(Supp. Fig. 1B)**. Further analysis of highly differentially expressed genes revealed only one overlapping gene (*CCDC190*) that was downregulated between flow and high-density conditions, and no genes were highly upregulated with both flow and high density conditions, indicating that the quiescence profiles of endothelial cells exposed to extended laminar flow or high density are largely distinct **(Supp. Fig. 1C-F)**. Further analysis revealed that expression of cell cycle activators was down-regulated regardless of quiescence induction stimulus, while cell cycle inhibitor expression showed an increased trend with high density but not with homeostatic flow **(Fig. 1D-E).** Thus, endothelial cells respond to homeostatic laminar flow exposure with a distinct quiescence transcriptional profile compared to high density-induced quiescence that includes differences in how cell cycle inhibitors are regulated, suggesting unique properties of flow-mediated endothelial cell quiescence.

### Shallow quiescence depth characterizes endothelial cells exposed to extended laminar flow

Quiescence depth, as defined by time to cell cycle reentry upon stimulus removal, varies in non-endothelial cells^22–24^. Because cell cycle inhibitor expression was not upregulated in endothelial cells exposed to laminar flow, we hypothesized that they are in a relatively shallow quiescence state. We assessed quiescence depth by measuring cell cycle reentry time after quiescence stimulus removal, via EdU labeling and pRB (phospho-retinoblastoma protein) expression **(Supp. Fig. 2A)**^37,38^. EdU labels cells in S-phase, and phosphorylation releases RB from E2F and permits progression through the G_1_-S checkpoint and proliferation^38,39^. Endothelial cells exposed to homeostatic laminar flow (Flow-M) or contact inhibition had significantly reduced EdU-labeled **(Fig. 2A-B, D-E)** and pRB+ cells **(Fig. 2A, C, D, F; Supp. Fig 2B-C)** at the end of the flow or contact inhibition period, consistent with previously published work^7,12,40^.

**Figure 2.**
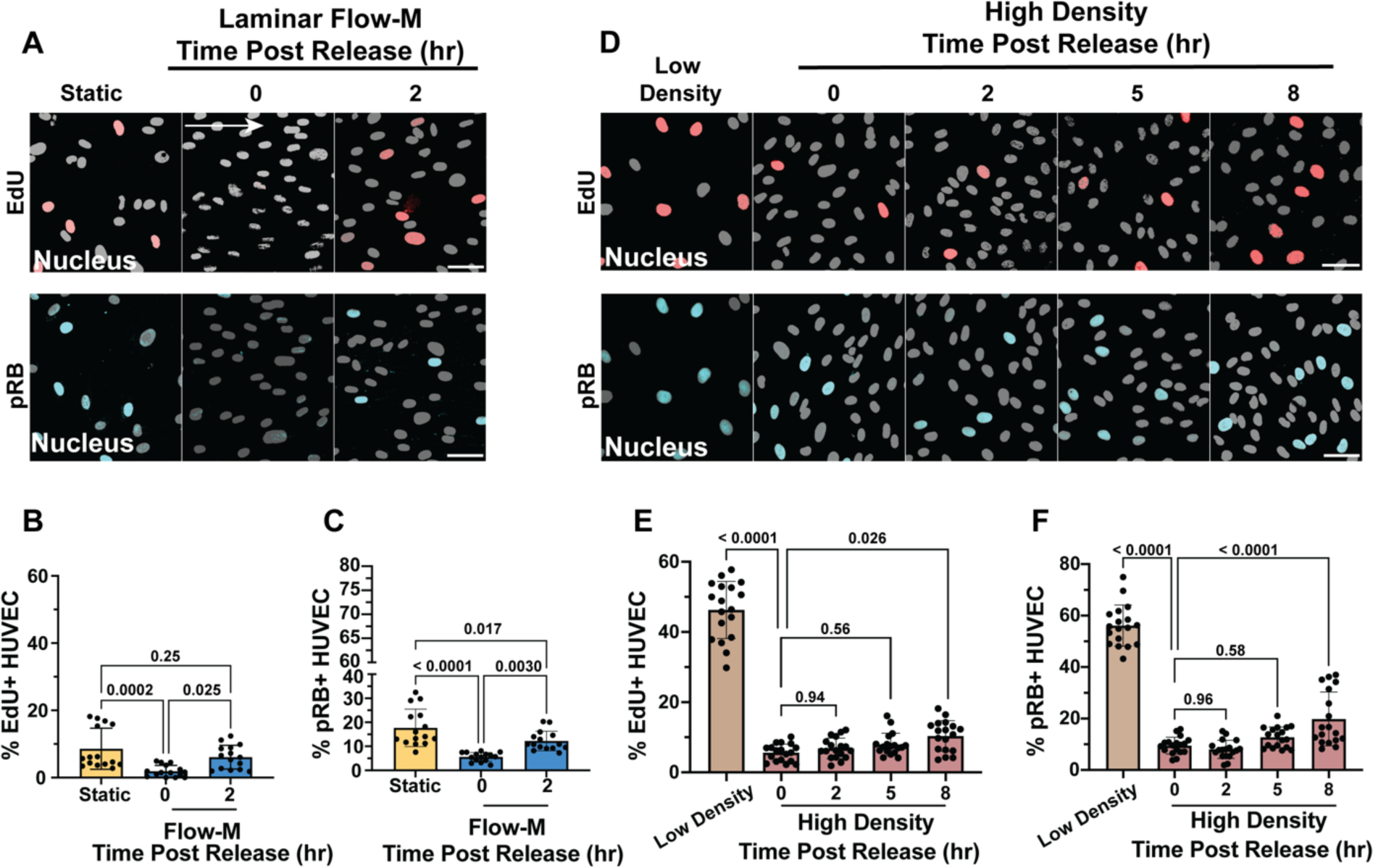
Endothelial cell flow maintenance quiescence stimulus leads to shallow quiescence depth. **(A)** Representative images of HUVEC under static (non-flow) or Flow-M (flow maintenance) conditions with EdU incorporation and fixation at indicated times post Flow-M release. Cultures stained for DAPI (white, nuclear mask) and EdU (red, S-phase) or pRB (blue, interphase). Scale bar, 50 µm. White arrow, flow direction. **(B)** Quantification of percent EdU+ cells with indicated conditions. n=3 replicates, 5 images per condition per replicate. **(C)** Quantification of percent pRB+ cells with indicated conditions. n=3 replicates, 5 images per condition per replicate. **(D)** Representative images of HUVEC under indicated density conditions with EdU incorporation and fixation at indicated times post high-density release. Cultures stained for DAPI (white, nuclear mask) and EdU (red, S-phase) or pRB (blue, interphase). Scale bar, 50 µm. **(E)** Quantification of percent EdU+ cells with indicated conditions. n=3 replicates, 6 images per condition per replicate. **(F)** Quantification of percent pRB+ cells with indicated conditions. n=3 replicates, 6 images per condition per replicate. Statistics, one-way ANOVA with Tukey’s multiple comparisons test.

HUVEC released from homeostatic flow (Flow-M) significantly increased EdU-labeled and pRB+ cells within 2h of flow cessation, while release from contact inhibition only showed significant increases in EdU-labeling and pRB reactivity 8h post-release **(Fig. 2A-F, Supp. Fig. 2B-C)**, and HAEC (human aortic endothelial cells) showed similar stimulus-dependent differences in quiescence depth **(Supp. Fig. 2D-G)**. Thus, endothelial cells exposed to homeostatic laminar flow (Flow-M) are less deeply quiescent compared to contact-inhibited cells, independent of endothelial subtype.

### Cell cycle inhibitor p27 expression correlates with quiescence depth and is required for flow-mediated quiescence

The cell cycle inhibitor p27 (*CDKN1B*) is often upregulated with quiescence in non-endothelial cells^41–43^, so we analyzed endothelial p27 expression and found a highly significant decrease in *CDKN1B* RNA expression and p27+ HUVEC after exposure to homeostatic Flow-M **(Fig. 3A-C; Supp. Fig 3A,C)**. In contrast, we confirmed that p27 RNA and protein levels significantly increased in contact inhibited HUVEC^40^ **(Fig. 3D-F; Supp. Fig. 3B-C)**, and HAEC also had reduced p27+ cells after exposure to homeostatic Flow-M **(Supp. Fig. 3D-E)**. Taken together, these findings show that p27 expression positively correlates with quiescence depth in endothelial cells, independent of endothelial subtype.

**Figure 3.**
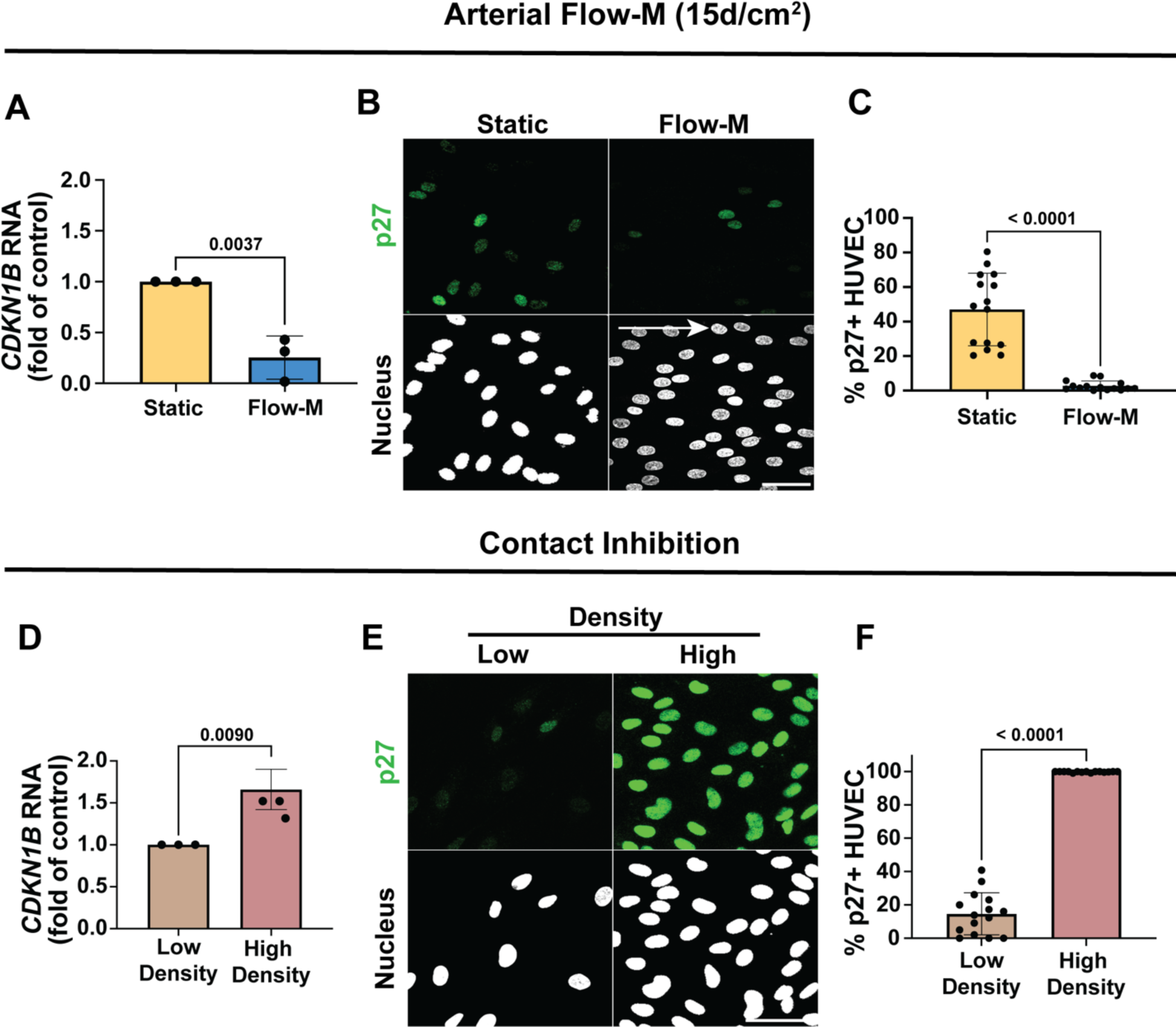
Cell cycle inhibitor p27 expression levels vary with endothelial quiescence stimulus. **(A)** RT-qPCR for *CDKN1B* levels under indicated conditions. n=3 replicates. **(B)** Representative images of HUVEC under indicated conditions stained for p27 (green) and DAPI (white, nuclear mask). Scale bar, 50 µm. White arrow, flow direction. **(C)** Quantification of HUVEC p27+ cells under indicated conditions. n=3 replicates, 5 images per condition per replicate. **(D)** RT-qPCR for *CDKN1B* levels under indicated conditions. n=3 replicates. **(E)** Representative images of HUVEC under indicated conditions stained for p27 (green) and DAPI (white, nuclear mask). Scale bar, 50 µm. **(F)** Quantification of HUVEC p27+ cells under indicated conditions. n=3 replicates, 5 images per condition per replicate. Statistics, student’s two-tailed *t*-test.

We interrogated p27 function in endothelial flow-mediated quiescence via siRNA knockdown (KD) and found that p27 depletion and exposure to homeostatic flow (Flow-M) significantly increased EdU-labeling and staining for Ki67, a marker of cells in S/G_2_/M^44^ over controls **(Fig. 4A-B; Supp. Fig. 3F-H)**. Transcriptional profiling of quiescent endothelial cells depleted for p27 under homeostatic flow or high density revealed little overlap in the profiles of genes up- or down-regulated between conditions (upregulated: 9.0% (12/133) flow and 8.6% (12/139) density shared genes; down-regulated 13.4% (18/134) flow and 0.4% (18/313) density shared genes) **(Supp. Fig. 4A)**. Further analysis of highly differentially expressed genes revealed no overlap between p27 depleted cells and controls in either condition, and gene ontology (GO) comparison revealed no shared terms in either condition **(Supp. Fig. 4B-I)**. These findings indicate that p27 is required for endothelial cell flow-mediated quiescence and affects transcriptional quiescence programs in a quiescence stimulus-dependent manner.

**Figure 4.**
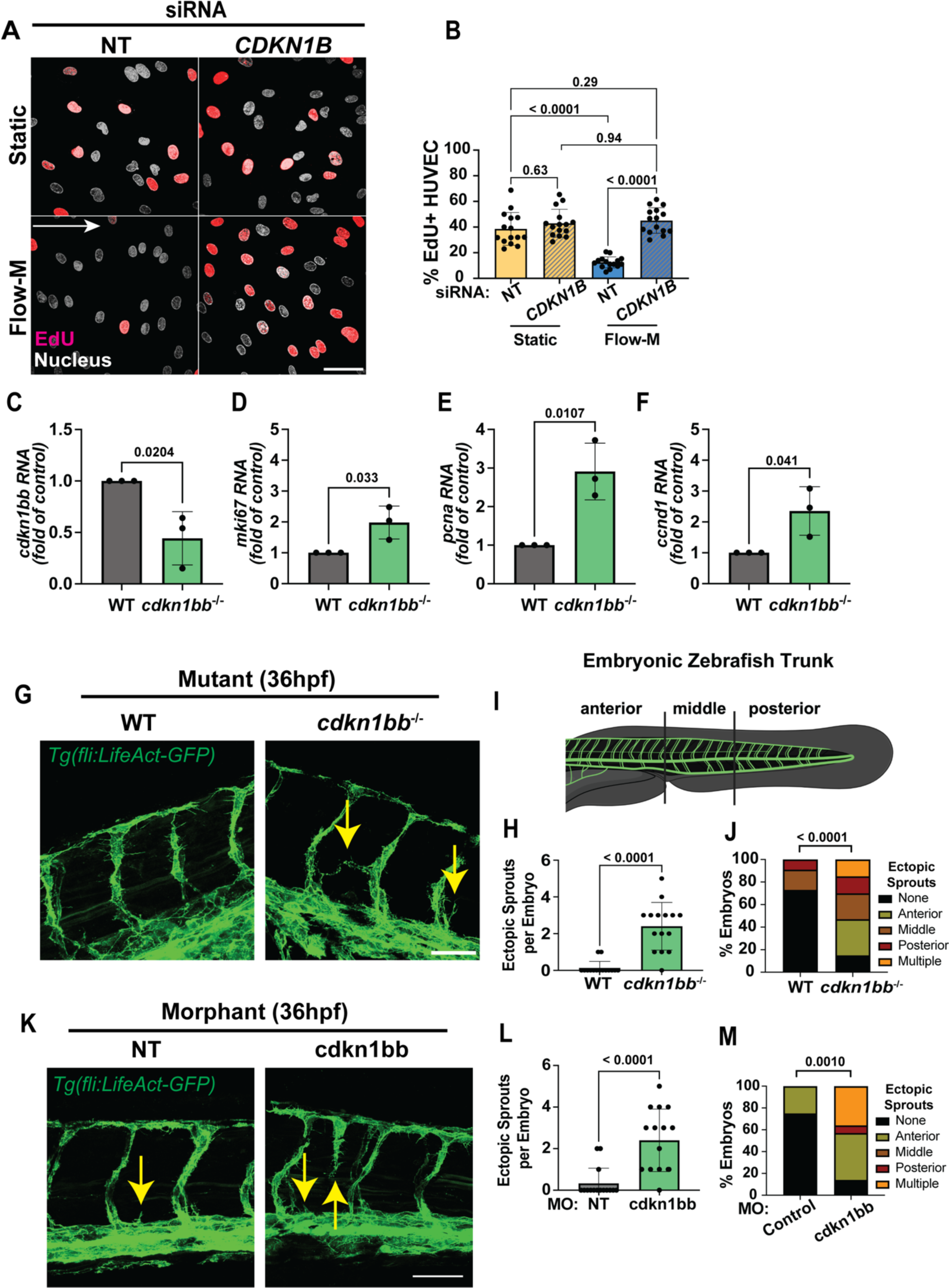
p27 establishes quiescence in endothelial cells and regulates cell cycle and vascular expansion *in vivo*. **(A)** Representative images of HUVEC with indicated siRNA treatments and conditions. after EdU incorporation (red, S-phase) and staining with DAPI (white, nuclear mask). Scale bar, 50 µm. White arrow, flow direction. **(B)** Quantification of EdU+ cells with indicated conditions. n=3 replicates, 5 images per condition per replicate. **(C-F)** RT-qPCR of FACs sorted 24 hpf zebrafish endothelial cells from *Tg(fli:LifeAct-GFP)* (green, endothelial cell marker) embryos of indicated genotypes for *cdkn1bb* **(C)**, *mki67* **(D)**, *pcna* **(E)**, and *ccnd1* **(F)** levels. n=3 replicates. **(G)** Representative images of 36hpf *Tg(fli:LifeAct-GFP)* (green, endothelial cell marker) embryos that were also WT or *cdkn1bb^-/-^*. Yellow arrows, ectopic sprouts. Scale bar, 50 µm. **(H)** Quantification of ectopic sprouts per embryo. n=3 replicates, 5 images per condition per replicate. **(I)** Diagram defining regions for ectopic sprout quantification in embryonic zebrafish. **(J)** Quantification of % ectopic sprouts in *Tg(fli:LifeAct-GFP)* (green, endothelial cell marker) embryos and of indicated genotypes per region. n=3 replicates, 5 images per condition per replicate. **(K)** Representative images of 36hpf *Tg(fli:LifeAct-GFP)* embryos injected with control (NT) or cdkn1bb MO at the one-cell stage. Yellow arrows, ectopic sprouts. Scale bar, 50 µm. **(L)** Quantification of ectopic sprouts per embryo. n=3 replicates, 5 images per condition per replicate. **(M)** Quantification of % ectopic sprouts in *Tg(fli:LifeAct-GFP)* (green, endothelial cell marker) and with indicated MO injection per region. n=3 replicates, 5 images per condition per replicate. Statistics, student’s two-tailed *t*-test (C-F, H, L), one-way ANOVA with Tukey’s multiple comparisons test (B), and Ξ^2^ test (J, M).

### p27 regulates endothelial quiescence parameters *in vivo*

We asked whether p27 affects vascular processes *in vivo* and hypothesized that p27 loss would deregulate the endothelial cell cycle and angiogenic expansion. We generated a *cdkn1bb* (zebrafish p27 gene) mutant zebrafish line and found significant increases in expression of cell cycle regulators Ki67 *(miKi67*), PCNA *(pcna)* and CyclinD1 *(ccnd1)* in enriched endothelial cell populations from mutant embryos that had reduced p27 (*cdkn1bb)* **(Fig. 4C-F)**. Analysis of vascular sprouting in *cdkn1bb^-/-^* fish revealed increased ectopic sprouts in the trunk vasculature that was phenocopied by morphant fish depleted for *cdkn1bb* **(Fig. 4G-H, K-L).** Regional quantification of ectopic sprouts along the anterior-posterior axis revealed that mutant or morphant embryos had more ectopic sprouts in the anterior position **(Fig. 4I-J, M)**, suggesting that effects of p27 loss are more associated with mature vascular regions that are likely transitioning to homeostasis. Thus, p27 is required for endothelial cell cycle regulation and proper sprouting *in vivo* and likely affects quiescence establishment.

### Flow-mediated quiescence depth is temporally regulated in endothelial cells

Since p27 is required for flow-mediated endothelial cell quiescence despite very low expression at 72h (Flow-M), we hypothesized that p27 levels temporally fluctuated with flow and found that p27+ cells significantly increased 16h after flow initiation, consistent with another study^45^, and then decreased over time to almost undetectable levels at 72h **(Fig. 5A-B, Supp. Fig. 5A).** Another CIP-KIP family cell cycle inhibitor, p21, showed a similar decrease in expression even earlier in the flow time course **(Fig. 5A,C, Supp. Fig. 5B)**, and Ki67+ cells decreased to almost undetectable levels over flow time **(Fig. 5A,D; Supp. Fig. 5C)**, consistent with the conclusion that endothelial cells leave the cell cycle and become quiescent by 16-24h of laminar flow, then subsequently down-regulate p27 levels while maintaining quiescence.

**Figure 5.**
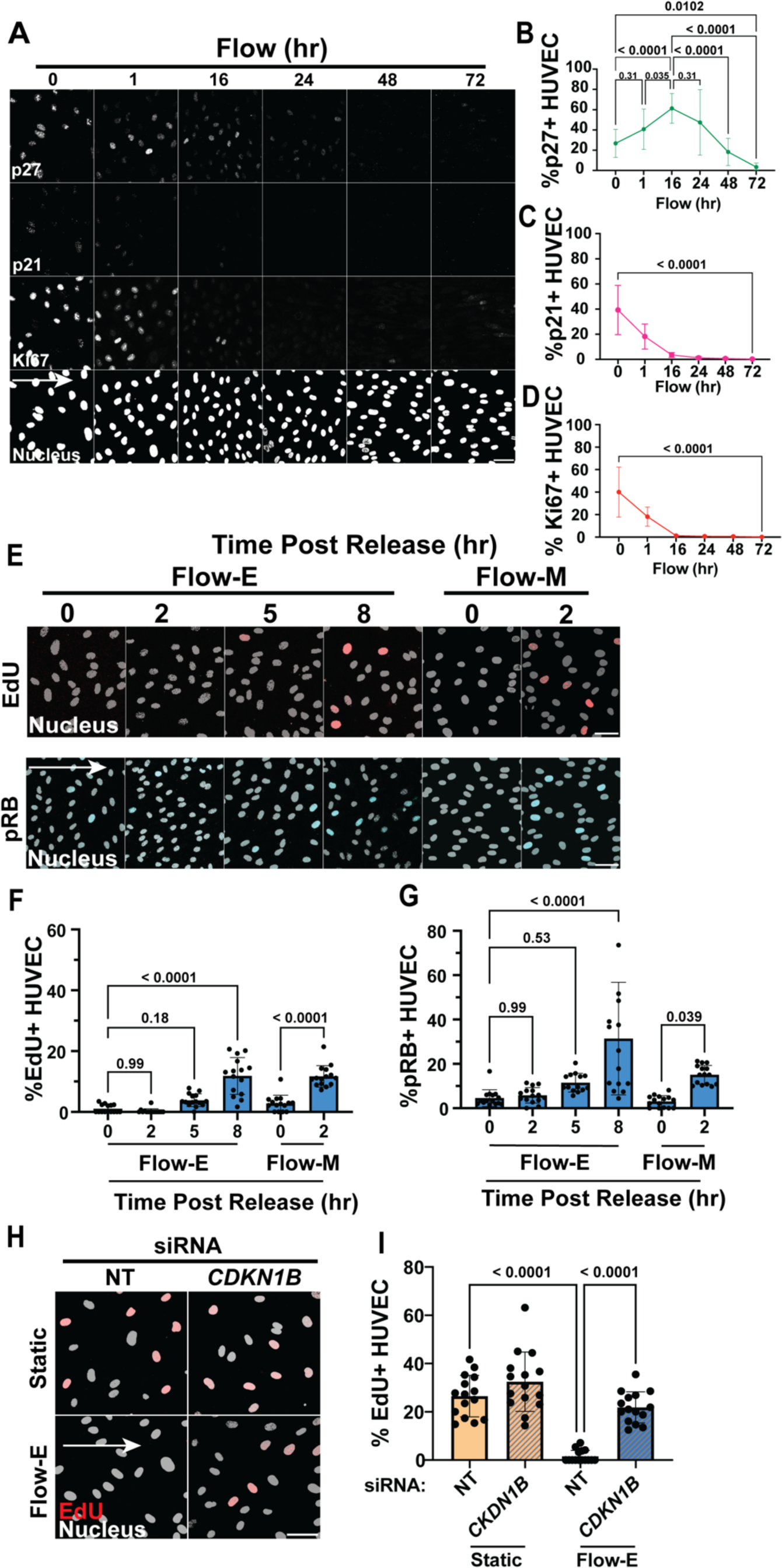
Endothelial quiescence depth varies with laminar flow exposure time and positively correlates with p27 expression levels. **(A)** Representative images of HUVEC with indicated conditions and stained for p27 (inhibitor), p21 (inhibitor), Ki67 (proliferation marker), and DAPI (nucleus) in white. Scale bar, 50 µm. White arrow, flow direction. **(B-D)** Quantification of percent p27+ (**B**), p21+ cells (**C**), or Ki67+ cells (**D**). n=3 replicates, 5 images per condition per replicate. **(E)** Representative images of HUVEC under indicted conditions and after EdU incorporation (red, S-phase) and stained for pRB (blue, interphase) and with DAPI (white, nuclear mask), Flow-E, flow establishment (15d/cm^2^, 16h); Flow-M, flow maintenance (15d/cm^2^, 72h). Scale bar, 50 µm. White arrow, flow direction. **(F)** Quantification of EdU+ cells under indicated conditions. n=3 replicates, 5 images per condition per replicate. **(G)** Quantification of pRB+ cells under indicated conditions. n=3 replicates, 5 images per condition per replicate. **(H)** Representative images of HUVEC with indicated siRNA treatments and conditions after EdU incorporation (red, S-phase) and staining with DAPI (white, nuclear mask). Scale bar, 50 µm. White arrow, flow direction. **(I)** Quantification of EdU+ cells with indicated conditions. n=3 replicates, 5 images per condition per replicate. Statistics, one-way ANOVA with Tukey’s multiple comparisons test.

We then asked whether endothelial cell quiescence depth was temporally regulated under laminar flow. After 16h of flow (flow establishment (Flow-E)), HUVEC only showed significant EdU-labeling and pRB reactivity 8h post-flow, similar to cells released from contact inhibition and longer than the 2h cell cycle reentry time exhibited by cells exposed to homeostatic flow (Flow-M) **(Fig. 5E-G; Supp. Fig. 5D)**. HAEC also displayed delayed cell cycle reentry post Flow-E, as measured by EdU+ labeling **(Supp. Fig. 5E-F)**. These findings correlate with temporal regulation of p27 levels over the flow period and indicate that endothelial cells initially respond to laminar flow with p27 upregulation and establishment of a relatively “deep” quiescence (Flow-E) that over time becomes shallower, along with reduced levels of cell cycle inhibitors p27 and p21. Consistent with this idea, endothelial cells were not aligned after 16h flow (Flow-E) but were aligned by 72h (Flow-M) under our flow conditions **(Supp. Fig. 5G-H).**

We further analyzed the relationship of endothelial quiescence and p27, via single-cell image analysis of endothelial cells exposed to Flow-E or high density and labeled with EdU and p27 reactivity at time points post-flow. Using a threshold for each label, no EdU+/p27+ cells were observed. With time post-flow, p27+/EdU-cells increased and then decreased while p27-/EdU+ cells increased, and with release from contact inhibition similar trends showed that p27+/EdU-cells were replaced by p27-/EdU+ cells over time **(Fig. Supp. 5I-J).** Functionally, endothelial cells with p27 depletion had elevated EdU labeling after 16h flow (Flow-E) **(Fig. 5H-I),** consistent with a previous report^45^. These findings indicate that endothelial cell quiescence depth positively correlates with p27 levels during the transition from deep to shallow quiescence, and that individual cells with low p27 expression become competent to reenter the cell cycle.

### Transcriptional targets of flow-mediated endothelial cell signaling regulate p27 levels under laminar flow

To better understand how endothelial cell p27 may regulate quiescence depth, we examined genes upregulated by laminar flow whose encoded proteins transcriptionally repress p27. Both Notch and BMP signaling are upregulated with laminar flow^2,9,10^, and *HES1* and *ID3* are downstream targets of these pathways that transcriptionally repress *CDKN1B* expression via direct and indirect promoter interactions^46,47^. Homeostatic flow (Flow-M) led to *HES1* and *ID3* RNA accumulation in HUVEC **(Supp. Fig. 6A-B)**, and depletion of either *HES1* or *ID3* RNA prevented the normal decrease in p27+ cells under homeostatic laminar flow (Flow-M) in both HUVEC and HAEC **(Fig. 6A-B; Supp. Fig. 6C-E)**, indicating that these repressors are required for flow-mediated reduction of p27 levels over time. Notch signaling is required for endothelial flow-mediated quiescence^45,48^, and consistent with this relationship, *NOTCH1* or *DLL4* depletion led to increased EdU+ cells after 16h flow (Flow-E) compared to control **(Supp. Fig. 6F-G)**.

**Figure 6.**
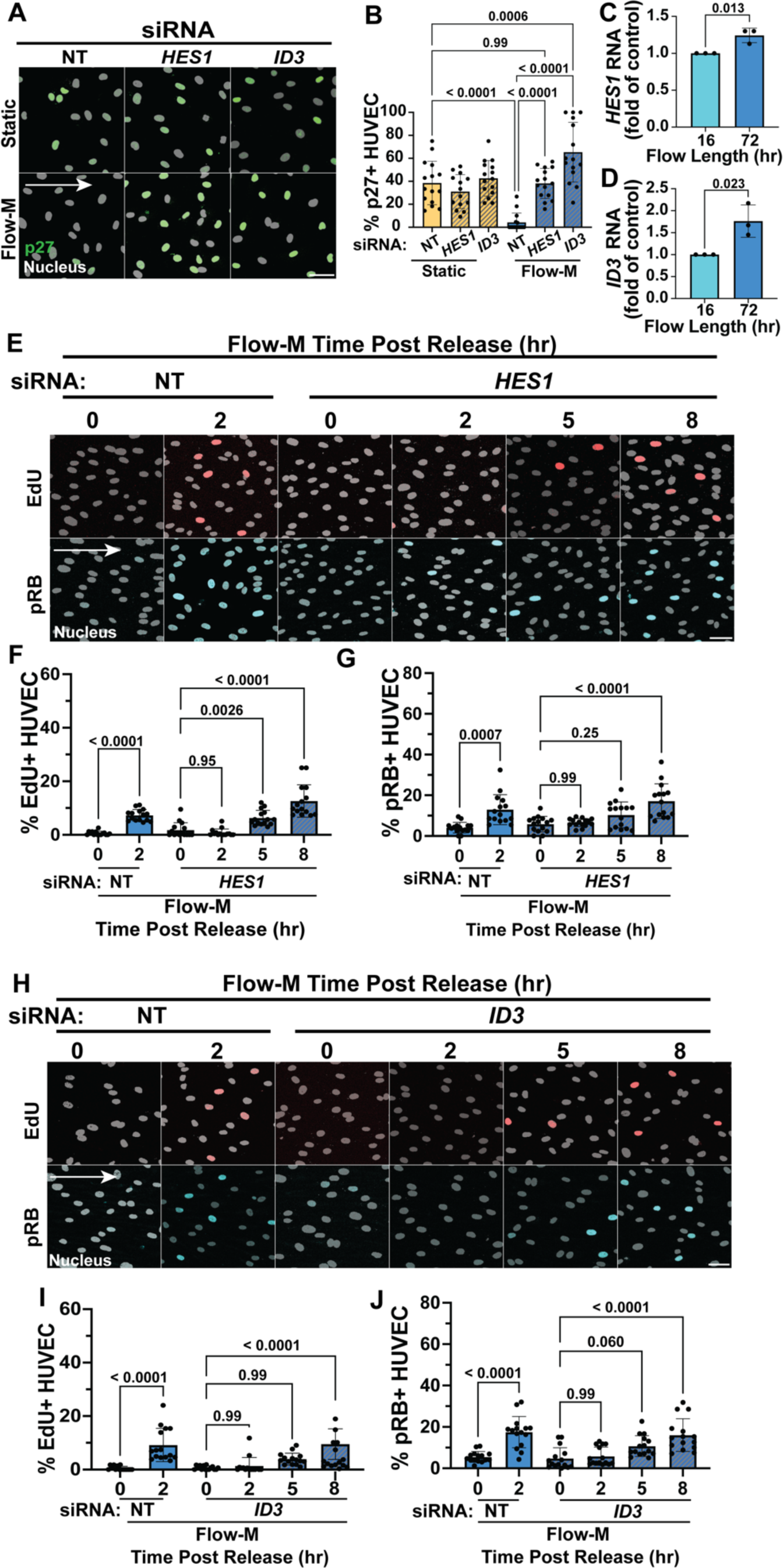
BMP and NOTCH regulated p27 repressors *HES1* and *ID3* regulate p27 levels and flow-mediated quiescence depth. **(A)** Representative images of HUVEC under indicated conditions and with indicated siRNA treatment. Cells stained for p27 (green) and DAPI (white, nuclear mask). Scale bar, 50 µm. White arrow, flow direction. **(B)** Quantification of p27+ cells under indicated conditions and treatments. n=3 replicates, 5 images per condition per replicate. **(C-D)** RT-qPCR for *HES1* **(C)** and *ID3* **(D)** levels under indicated conditions. n=3 replicates. **(E)** Representative images of HUVEC under indicated conditions and siRNA treatments. Cells were labeled with EdU (red, S-phase) and stained for pRB (blue, interphase) and DAPI (white, nuclear mask). Scale bar, 50 µm. White arrow, flow direction. **(F)** Quantification of EdU+ cells with indicated conditions. n=3 replicates, 5 images per condition per replicate. **(G)** Quantification of pRB+ cells with indicated conditions. n=3 replicates, 5 images per condition per replicate. **(H)** Representative images of HUVEC under indicated conditions and siRNA treatments. Cells were labeled with EdU (red, S-phase) and stained for pRB (blue, interphase) and DAPI (white, nuclear mask). Scale bar, 50 µm. White arrow, flow direction. **(I)** Quantification of EdU+ cells under indicated conditions. n=3 replicates, 5 images per condition per replicate. **(J)** Quantification of pRB+ cells under indicated conditions. n=3 replicates, 5 images per condition per replicate. Statistics, student’s two-tailed *t*-test (C-D) and one-way ANOVA with Tukey’s multiple comparisons test (B, F-G, I-J).

We hypothesized that *HES1* and *ID3* mediate the temporal changes to endothelial cell quiescence depth and predicted that *HES1* or *ID3* depletion would prevent transition from deep to shallow quiescence. *HES1* and *ID3* expression were significantly increased in cells exposed to Flow-M vs. Flow-E **(Fig. 6C-D)**, and either *HES1* or *ID3* depletion significantly extended the time required for significant increases in EdU+ and pRB+ endothelial cells after Flow-M **(Fig. 6E-J, Supp. Fig. 6H-I),** suggesting that the normal temporal change to shallow quiescence was prevented when these p27 repressors were depleted. These findings are consistent with the idea that endothelial quiescence depth is temporally regulated from deep to shallow over time in conjunction with laminar flow-mediated changes, and that p27 regulates the transition downstream of the transcriptional repressors *HES1* and *ID3*.

### Endothelial cells exposed to venous laminar flow maintain deep quiescence

We asked whether temporal flow-mediated quiescence depth changes depended on the flow vector (shear stress) magnitude, and we predicted that a lower magnitude of flow characteristic of venous flow would be insufficient to repress p27 levels over time. In contrast to arterial laminar flow (15d/cm^2^), endothelial cell exposure to venous laminar flow (5d/cm^2^) for 72h (Flow-MV) did not decrease p27+ cells and had increased p27 RNA expression over flow time (Flow-EV) **(Fig. 7A-C)**. Consistent with p27 levels, endothelial cells exposed to venous flow required extended time (8h) to significantly increase EdU+ cells associated with cell cycle reentry **(Fig. 7D-G)** at both 16h (Flow-EV) and 72h (Flow-MV), indicating that temporal quiescence depth regulation depends on the magnitude of the flow stimulus.

**Figure 7.**
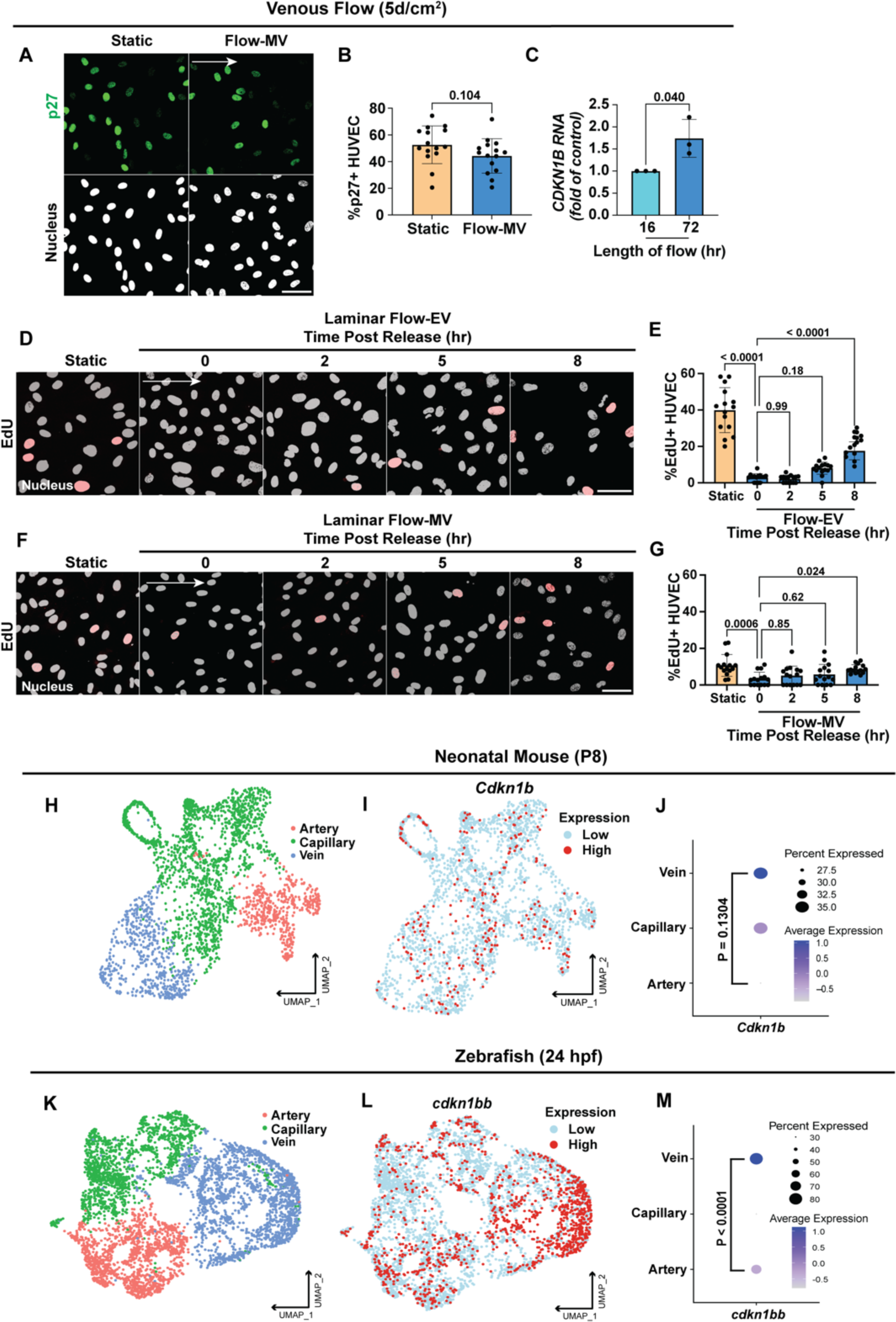
Endothelial cells establish and maintain deep quiescence under venous flow and express elevated p27 levels *in vivo*. **(A)** Representative images of HUVEC under indicated conditions (Flow-MV (5d/cm^2^, 72h)) stained for p27 (green) and DAPI (white, nuclear mask). Scale bar, 50 µm. White arrow, flow direction. **(B)** Quantification of HUVEC p27+ cells under indicated conditions, n=3 replicates, 5 images per condition per replicate. **(C)** RT-qPCR for *CDKN1B* levels in indicated conditions. n=3 replicates. **(D)** Representative images of HUVEC under static (non-flow) or Flow-EV (5d/cm^2^, 16h) conditions with EdU incorporation and fixation at indicated times post Flow-EV release. Cells stained for DAPI (white, nuclear mask) and EdU (red, S-phase), Scale bar, 50 µm. White arrow, flow direction. **(E)** Quantification of EdU+ cells with indicated conditions. n=3 replicates, 5 images per condition per replicate. **(F)** Representative images of HUVEC under static (non-flow) or Flow-MV conditions (5d/cm^2^, 72h) with EdU incorporation and fixation at indicated times post Flow-MV release. Cells stained for DAPI (white, nuclear mask) and EdU (red, S-phase), Scale bar, 50 µm. White arrow, flow direction. **(G)** Quantification of EdU+ cells with indicated conditions. n=3 replicates, 5 images per condition per replicate. **(H)** UMAP grouping of artery, vein, and capillary endothelial cell clusters enriched from neonatal mouse ear skin. **(I)** UMAP overlaid with *Cdkn1b (p27)* expression. **(J)** Dot plot of *Cdkn1b* expression by endothelial sub-type. **(K)** UMAP grouping of endothelial artery, vein, and capillary clusters reanalyzed from 24hpf embryonic zebrafish. **(L)** UMAP overlaid with *cdkn1bb (p27)* expression. **(M)** Dot plot of *Cdkn1b* expression by endothelial sub-type. Statistics, student’s two-tailed *t*-test (B-C, J-M) and one-way ANOVA with Tukey’s multiple comparisons test (E, G).

Since quiescence depth is linked to p27 levels in flow-exposed cultured cells, and since flow manipulations are imprecise *in vivo*, we used p27 levels as a proxy for quiescence depth *in vivo* and hypothesized that venous vs. arterial comparisons would reveal elevated venous p27 levels. Two endothelial scRNAseq datasets, one from neonatal mouse skin generated by our lab and one from 24hpf zebrafish generated by the Sumanas lab^49^, were subjected to UMAP analysis to define artery/vein/capillary clusters **(Fig. 7H, K)**, and we found that *Cdkn1b* (mouse p27 gene) levels trended higher in venous clusters (p = 0.1304) compared to arterial clusters **(Fig. 7I-J)** and were highly significantly increased in zebrafish venous clusters (**Fig 7L-M**, p=0.0001). These findings show that p27 levels correlate with flow magnitude *in vivo* and suggest that quiescence depth differs with flow magnitude in arteries vs. veins **(Fig. 8)**.

**Figure 8.**
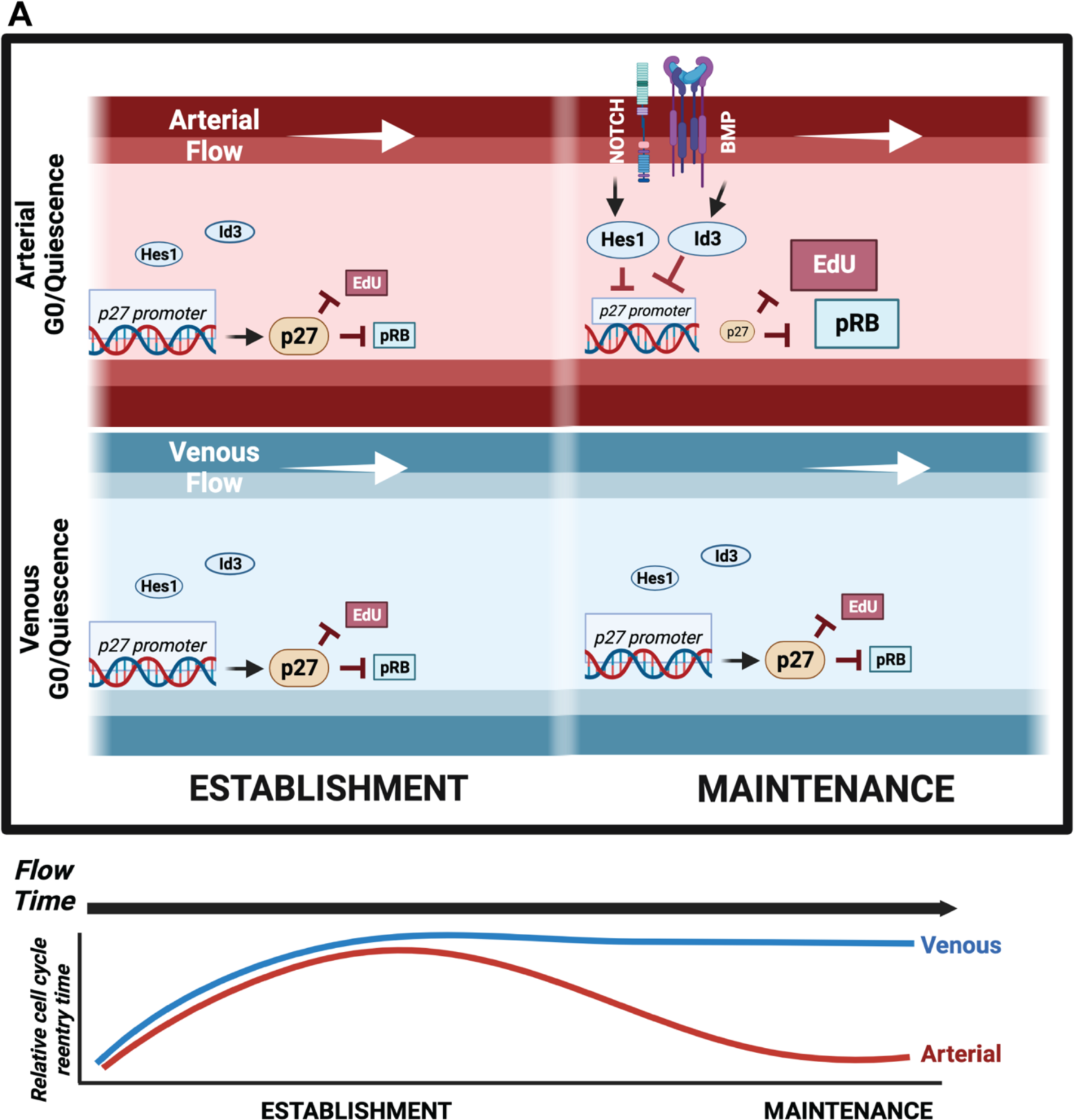
Model of endothelial cell flow-mediated quiescence depth. Proposed model for dynamic regulation of endothelial cell flow-mediated quiescence depth with flow exposure time and magnitude. In the first 16h (Flow-E, arterial flow establishment) of laminar flow, a deep quiescence is established accompanied by high levels of cell cycle inhibitor p27, independent of flow magnitude. With time under arterial flow (Flow-M, flow maintenance), *HES1* and *ID3* transcription factors are upregulated downstream of flow-mediated Notch and BMP signaling, and they repress p27 transcription leading to a shallow quiescence depth. Deep quiescence and high p27 levels characterize venous flow establishment at 16h (Flow-EV, venous flow establishment), and this deep quiescence perdures with time under venous flow (Flow-MV).

## DISCUSSION

Flow-mediated endothelial cell quiescence is important for vascular homeostasis and proper barrier function, but how it is set up and maintained is not well-understood. Here we show that endothelial cell quiescence varies with stimulus and flow magnitude and is temporally regulated under flow. We also confirm a functional requirement for the cell cycle inhibitor p27 and find that expression positively correlates with quiescence depth in cells and varies with flow magnitude *in vivo*. Temporal quiescence depth changes under arterial flow require flow-regulated p27 repressors that are targets of flow-mediated pathways such as Notch, suggesting complex regulation of endothelial cell quiescence depth and a model of quiescence regulation **(Fig. 8)**, and they reveal new critical control points for vascular homeostasis^50–52^.

A newly generated endothelial quiescence score tracked well with endothelial cells made quiescent via contact inhibition or homeostatic laminar flow (Flow-M), and application of a published quiescence score developed using different cells and criteria^34^ revealed similar trends, indicating that some aspects of vascular quiescence are shared among cell types and conditions. However, further transcriptome analysis revealed stimulus-dependent differences in quiescent endothelial cells, and only contact inhibition broadly stimulated expression of cell cycle inhibitor genes. Further analysis revealed that most differentially regulated genes were unique to the quiescence stimulus, and comparison of highly regulated genes supported that transcriptional profiles are largely unique to the stimulus used to induce quiescence in endothelial cells, similar to patterns in fibroblasts^18^.

Endothelial cells under homeostatic arterial flow (Flow-M) reentered the cell cycle shortly after flow cessation while contact released cells took significantly longer, indicating distinct quiescence depths associated with quiescence stimulus. Temporal analysis revealed that under Flow-M, that simulates arterial levels of shear stress, endothelial cells first set up a deep quiescence that becomes shallow over flow time, while under Flow-MV, that simulates venous flow, deep quiescence was established in the same time frame but not changed over flow time. This provocative finding suggests that deep quiescence may be important in venous vascular beds that are more poised to proliferate^53,54^, to prevent inappropriate cell cycle reentry absent an angiogenic stimulus. It is also in line with a recent study showing that injury-induced collateral vessel expansion originates from arterial endothelial cells^55^, and that the inverse correlation of quiescence depth at homeostasis and flow magnitude is linked to interstitial flow where elevated levels are associated with shallower quiescence in fibroblasts^56^. Taken together, these findings suggest complex regulation of quiescence depth relative to flow stimulus.

Expression of the cell cycle inhibitor p27 positively correlated with quiescence depth in all scenarios, including between homeostatic Flow-M (p27 levels low) vs. contact inhibition (p27 levels high)^40^; between arterial Flow-E (establishment) vs. Flow-M (maintenance) flow times; and between flow magnitude, with lower magnitude venous-type flow associated with elevated p27 levels and deep quiescence across time. This strong correlation was mirrored *in vivo*, as p27 expression was elevated in endothelial cells identified as venous vs. arterial in neonatal mouse ears and zebrafish embryos. We confirmed that p27 is required to establish quiescence in cultured endothelial cells^45^ independent of flow magnitude, and we found that p27 loss leads to cell cycle mis-regulation and expanded sprouting *in vivo*, indicating that p27 regulates quiescence establishment and depth in physiological angiogenesis.

The temporal fluctuations in quiescence depth and p27 levels under arterial flow link to *HES1* and *ID3*, flow-regulated^9,10^ transcriptional repressors of p27^57–59^ downstream of Notch and BMP signaling, respectively. Both repressors are functionally required non-redundantly to dampen p27 expression as endothelial cells move from quiescence establishment to maintenance and vascular homeostasis, and their expression increases with flow time as cells transition from establishment to maintenance. Quiescence depth under arterial flow is more profound in the absence of either repressor, suggesting that increased Notch and BMP signaling during establishment, and subsequent upregulation of these repressors leads to reduced p27 expression levels and a change from a deep to shallow endothelial quiescence depth **(Fig. 8).** This model suggests that in addition to p27-mediated cell cycle regulation in early vascular development to promote arterio-venous differentiation^45,60^, p27 is important for subsequent modulation of endothelial quiescence depth.

What might be an advantage to endothelial cells in a shallow quiescence state under arterial flow maintenance (homeostasis) conditions? One possibility is protection from premature permanent arrest, called senescence^61^. Quiescent fibroblasts eventually undergo senescence^62^, and fibroblast *HES1* expression prevents premature senescence^63^. Senescent arterial endothelial cells often express a senescence-associated secretory phenotype (SASP)^64^ characterized by inflammatory cytokine expression that contributes to endothelial dysfunction, leading to cardiovascular disease^65^. Thus, arterial endothelial cells in a shallow quiescence state may be protected from inappropriate senescence, while the deeper quiescence of venous endothelial cells may help prevent inappropriate cell cycle reentry. Thus, our findings that endothelial flow-mediated quiescence regulation is linked to p27 suggest potential new therapeutic targets for vascular dysfunction.

## ACKNOWLEDGMENTS

We thank members of the Bautch and Cook labs for their support and feedback, the UNC High Throughput Sequencing Core for technical help, and the UNC Flow Cytometry Core for FACs sorting. We thank BioRender for figure model generation.

## AUTHOR CONTRIBUTIONS

Natalie T Tanke (NTT), Jeanette Gowen Cook (JGC), Victoria L Bautch (VLB) conceptualized the work; NTT, Ziqing Liu (ZL), Michaelanthony T Gore (MTG), Pauline Bougaran (PB), Allison Marvin (AM), Mary B Linares (MBL), Arya Sharma (AS), Morgan Oatley (MO), Tianji Yu (TY), Kaitlyn Quigley (KQ), and Sarah Vest (SV) performed and analyzed experiments. NTT and VLB wrote and edited the manuscript; VLB and JGC provided study supervision and oversight.

## FUNDING

This work was supported by grants from the National Institutes of Health (R35 HL139950 and GM129074 to VLB), the MiBioX T32 Training Grant (T32 GM 119999-03), a Ruth L. Kirschstein Predoctoral Fellowship (1F31HL156527-01 to NTT), and an American Heart Association Postdoctoral Fellowship (AHA829371 to MO).

## DECLARATION OF INTERESTS

none

## Nonstandard Abbreviations and Acronyms

HUVEC: Human Umbilical Vein Endothelial Cells
HAEC: Human Aortic Endothelial Cells
Flow-M: Flow maintenance 72 hr 15d/cm^2^
Flow-E: Flow establishment 16 hr 15d/cm^2^
Flow-EV: Flow establishment venous 16h, 5d/cm^2^
Flow-MV: Flow maintenance, venous 16h, 5d/cm^2^
siRNA: Small interfering RNA
KD: Knockdown A.U.: Arbitrary unit
pRB: Phospho retinoblastoma protein
EdU: 5-Ethynyl-2’-deoxyuridine
*HES1*: Hes Family BHLH Transcription Factor 1
*ID3*: Inhibitor of DNA Binding 3

## SUPPLEMENTARY FIGURE LEGENDS

**Supplemental Figure 1. Endothelial cell quiescence transcriptional profiles are stimulus-dependent.**

**(A)** Quantification using an epithelial quiescence score^34^ on HUVEC bulk RNAseq dataset under different conditions. n=3 replicates. Flow-M, flow maintenance (15d/cm^2^, 72h). **(B)** Venn diagrams showing bulk RNA seq analysis of genes up- and down-regulated in HUVEC under indicated conditions. **(C-F)** Heatmaps showing relative expression levels of differentially regulated genes (top 50 by fold change and p-value) in bulk RNA seq from HUVEC under indicated conditions. Statistics, one-way ANOVA with Tukey’s multiple comparisons test.

**Supplemental Figure 2. Quiescence depth is replicated in HAEC (human arterial endothelial cells).**

**(A)** Schematic showing areas of scratch wound used for imaging post-scratch for contact inhibition quiescence depth experiments. **(B)** Quantification of HUVEC pRB nuclear fluorescence intensity under indicated conditions. n=3 replicates, 5 images averaged per condition per replicate. **(C)** Quantification of HUVEC nuclear fluorescence intensity of pRB in low vs. high density release timepoints. n=3 replicates, 5 averaged images per condition per replicate. **(D)** Representative images of HAEC under static (non-flow) or Flow-M (flow maintenance) conditions with EdU incorporation and fixation at indicated times post Flow-M release. Cells stained for DAPI (white, nuclear mask) and EdU (red, S-phase). Scale bar, 50 µm. White arrow, flow direction. **(E)** Quantification of percent EdU+ cells with indicated conditions. n=3 replicates, 5 images per condition per replicate. **(F)** Representative images of HAEC under indicated density conditions with EdU incorporation and fixation at indicated times post density release. Cells stained for DAPI (white, nuclear mask) and EdU (red, S-phase). Scale bar, 50 µm. **(G)** Quantification of percent EdU+ cells with indicated conditions. n=3 replicates, 5 images per condition per replicate. Statistics, one-way ANOVA with Tukey’s multiple comparisons test.

**Supplemental Figure 3. Cell cycle inhibitor p27 expression differs with quiescence stimulus.**

**(A)** Quantification of p27 nuclear fluorescence intensity in indicated conditions. n=3 replicates, 5 images averaged per condition per replicate. **(B)** Quantification of p27 nuclear fluorescence intensity in indicated conditions. n=3 replicates, 5 images averaged per condition per replicate. **(C)** Western blot of p27 expression under indicated conditions, and with indicated antibodies. p27 Ab #1 (Cell Signaling) and p27 Ab #2 (Santa Cruz). **(D)** Representative images of HAEC under indicated conditions stained for p27 (green) and DAPI (white, nuclear mask). Scale bar, 50 µm. White arrow, flow direction. **(E)** Quantification of HAEC p27+ cells under indicated conditions, n=3 replicates, 5 images per condition per replicate. **(F)** Representative images of HUVEC with indicated siRNA and treatments. Endothelial cells stained with Ki67 (yellow, proliferation marker) and DAPI (gray, nucleus mask). Scale bar 50 µm. White arrow, flow direction. **(G)** Quantification of percent Ki67+ cells. n=3 replicates, 5 images per condition per replicate. **(H)** Quantification of nuclear fluorescence intensity of Ki67. n=3 replicates, 5 averaged images per condition per replicate. Statistics, student’s two-tailed *t*-test (A-B, E) and one-way ANOVA with Tukey’s multiple comparisons test (G-H).

**Supplemental Figure 4. p27 depletion leads to distinct transcriptional changes dependent on quiescence stimulus.**

**(A)** Venn diagrams showing overlap of HUVEC genes differentially regulated in indicated conditions and for *CDKN1B* KD compared to NT. **(B-E)** Heatmaps showing relative expression levels of genes differentially regulated (top 50 by fold change and p-value) in response to indicated conditions. **(F-I)** GO analysis performed on differentially expressed genes from bulk RNA-seq data comparing indicated conditions/treatments. Representative biological processes GO terms significantly enriched (P adjusted < 0.1) in differentially regulated genes are shown.

**Supplemental Figure 5. Cell cycle inhibitor and Ki67 expression intensity varies with flow and correlates with flow-mediated endothelial cell alignment.**

**(A)** Quantification of p27 nuclear fluorescence intensity under indicated conditions. n=3 replicates, 5 averaged images per condition per replicate. **(B)** Quantification of p21 nuclear fluorescence intensity under indicated conditions. n=3 replicates, 5 averaged images per condition per replicate. **(C)** Quantification of Ki67 nuclear fluorescence intensity under indicated conditions. n=3 replicates, 5 averaged images per condition per replicate. **(D)** Quantification of pRB nuclear fluorescence intensity under indicated conditions. n=3 replicates, 5 averaged images per condition per replicate. **(E)** Representative images of HAEC under static (non-flow) or Flow-E conditions with EdU incorporation and fixation at indicated times post Flow-E release. Cells stained for DAPI (white, nuclear mask) and EdU (red, S-phase), Scale bar, 50 µm. White arrow, flow direction. **(F)** Quantification of EdU+ cells under indicated conditions. n=3 replicates, 5 images per condition per replicate. **(G)** Representative images of HUVEC stained with VE-cadherin (white, junction marker) and DAPI (blue, nucleus) under indicated conditions. Scale bar, 50 µm. White arrow, flow direction. **(H)** Cell axis ratio quantification under indicated conditions. n=3 replicates, 5 images per condition per replicate. **(I)** % HUVEC under Flow-E release with p27-/EdU-, p27+/EdU-, and p27-/EdU+ incorporation. **(J)** % HUVEC under high density release with p27-/EdU-, p27+/EdU-, and p27-/EdU+ incorporation. Statistics, one-way ANOVA with Tukey’s multiple comparisons test (A-D, F, H) and Ξ^2^ test (I-J).

**Supplemental Figure 6. Nuclear fluorescence intensity of markers and changes with HES1 or ID3 depletion.**

**(A)** RT-qPCR for *HES1* expression under indicated conditions. n=3 replicates. **(B)** RT-qPCR for *ID3* expression under indicated conditions, n=3 replicates. **(C)** Quantification of p27 nuclear fluorescence intensity with indicated siRNA treatments and conditions. n=3 replicates, 5 averaged images per condition per replicate. **D)** Representative images of HAEC under indicated conditions and with indicated siRNA treatment. Cells stained for p27 (green) and DAPI (white, nuclear mask). Scale bar, 50 µm. White arrow, flow direction. **(E)** Quantification of p27+ cells under indicated conditions and treatments. n=3 replicates, 5 images per condition per replicate. **(F)** Representative images of HUVEC under indicated conditions and siRNA treatments. Cells were labeled with EdU (red, S-phase) and stained for DAPI (white, nuclear mask). Scale bar, 50 µm. White arrow, flow direction. **(G)** Quantification of EdU+ cells with indicated conditions. n=3 replicates, 5 images per condition per replicate. **(H)** Quantification of pRB nuclear fluorescence intensity under indicated siRNA treatments and conditions. n=3 replicates, 5 averaged images per condition per replicate. **(I)** Quantification of pRB nuclear fluorescence intensity under indicated siRNA treatments and conditions. n=3 replicates, 5 averaged images per condition per replicate. Statistics, student’s two tailed t-test (A-B) and one-way ANOVA with Tukey’s multiple comparisons test (C, E, G-I).

## Notes

### Competing Interest Statement

The authors have declared no competing interest.

### Summary of Updates

This version of the manuscript has been revised to incorporate new in vivo data and different fluidic data, as reflected in the text and images.

