## Supplemental Figures and Tables for "Endothelial cell flow-mediated quiescence is temporally regulated and utilizes the cell cycle inhibitor p27"

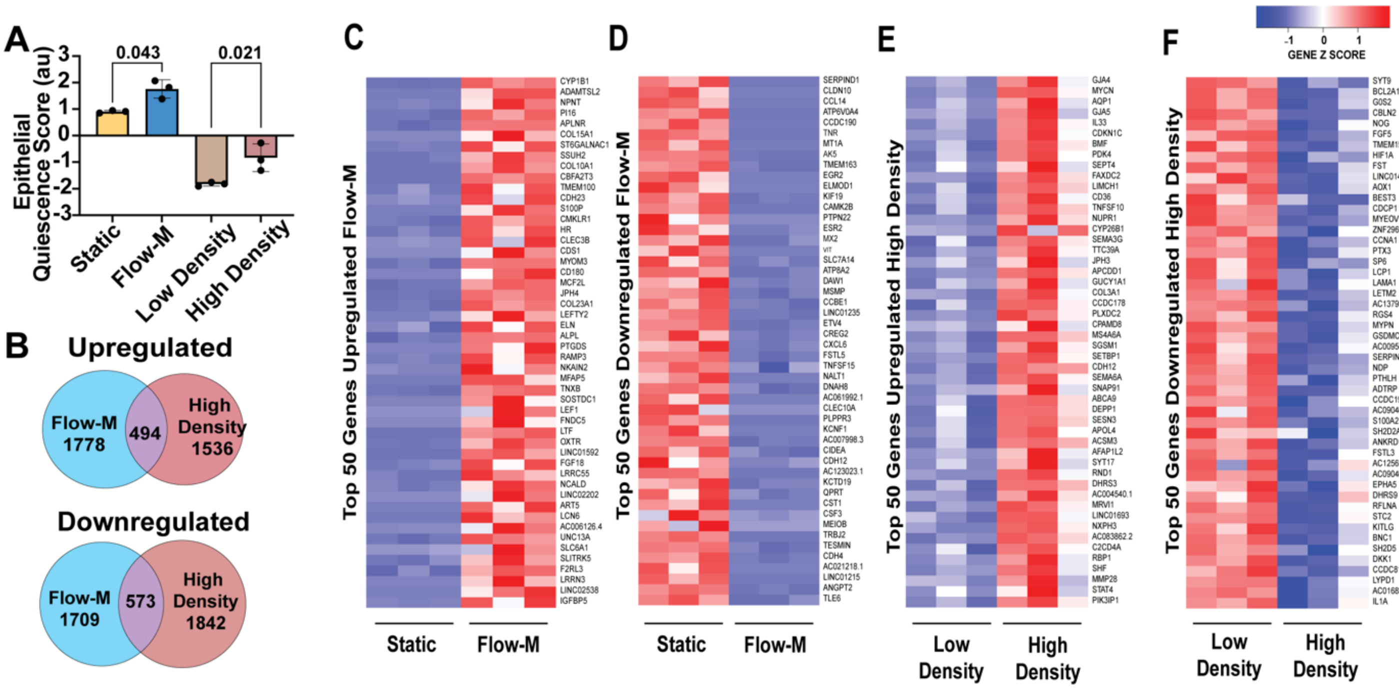

**Supplemental Figure 1. Endothelial cell quiescence transcriptional profiles are stimulus-dependent.**

**(A)** Quantification using an epithelial quiescence score[53] on HUVEC bulk RNAseq dataset under different conditions.  n=3 replicates. Flow-M, flow maintenance (15d/cm$^2$, 72h). **(B)** Venn diagrams showing bulk RNA seq analysis of genes up- and down-regulated in HUVEC under indicated conditions. **(C-F)** Heatmaps showing relative expression levels of differentially regulated genes (top 50 by fold change and p-value) in bulk RNA seq from HUVEC under indicated conditions. Statistics, one-way ANOVA with Tukey's multiple comparisons test.

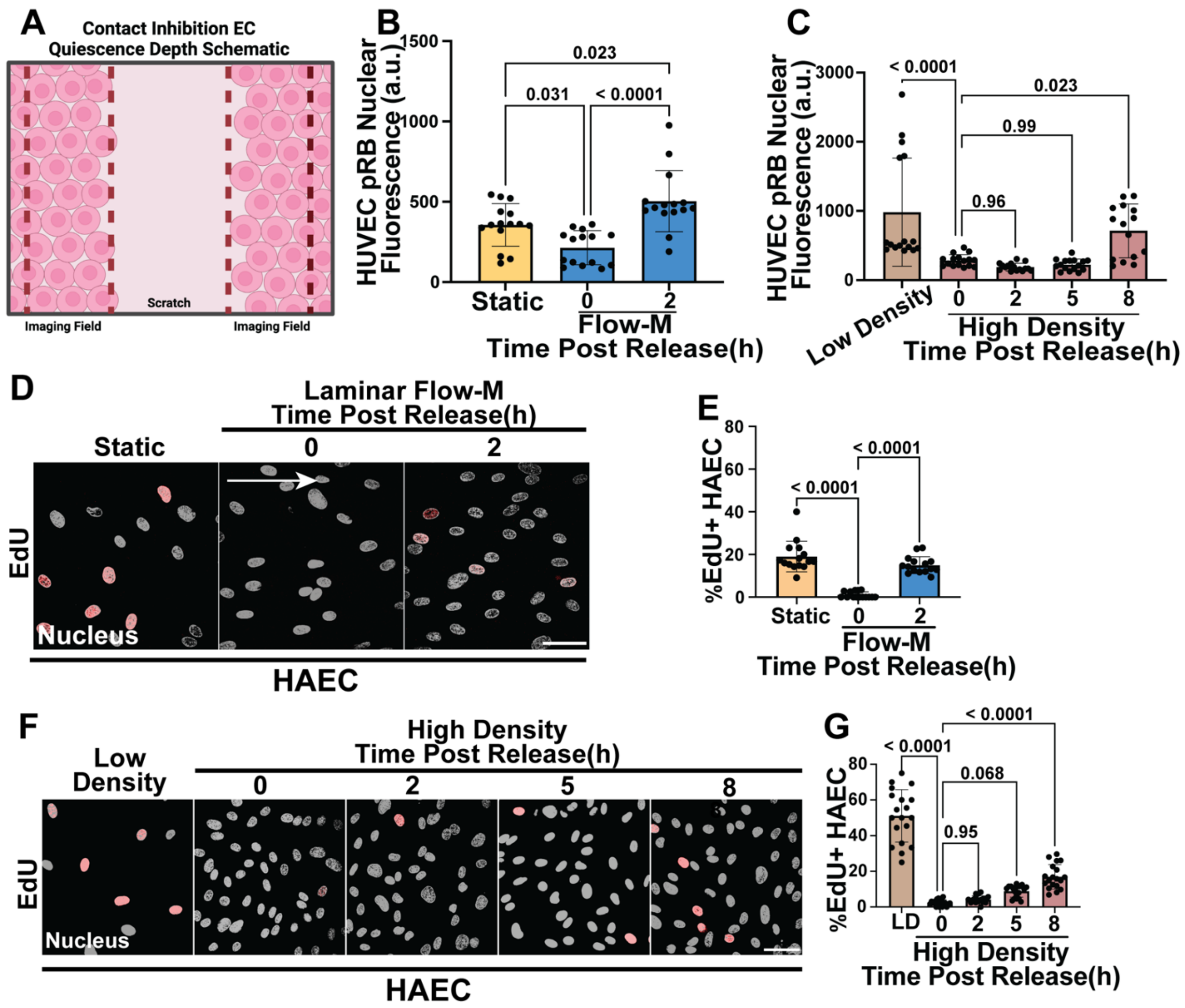


**Supplemental Figure 2. Quiescence depth is replicated in HAEC (human arterial endothelial cells).**

(A) Schematic showing areas of scratch wound used for imaging post-scratch for contact inhibition quiescence depth experiments. **(B)** Quantification of HUVEC pRB nuclear fluorescence intensity under indicated conditions. n=3 replicates, 5 images averaged per condition per replicate. **(C)** Quantification of HUVEC nuclear fluorescence intensity of pRB in low vs. high density release timepoints. n=3 replicates, 5 averaged images per condition per replicate. **(D)** Representative images of HAEC under static (non-flow) or Flow-M (flow maintenance) conditions with EdU incorporation and fixation at indicated times post Flow-M release. Cells stained for DAPI (white, nuclear mask) and EdU (red, S-phase). Scale bar, 50 µm. White arrow, flow direction. **(E)** Quantification of percent EdU+ cells with indicated conditions. n=3 replicates, 5 images per condition per replicate. **(F)** Representative images of HAEC under indicated density conditions with EdU incorporation and fixation at indicated times post density release. Cells stained for DAPI (white, nuclear mask) and EdU (red, S-phase). Scale bar, 50 µm. **(G)** Quantification of percent EdU+ cells with indicated conditions. n=3 replicates, 5 images per condition per replicate. Statistics, one-way ANOVA with Tukey's multiple comparisons test.

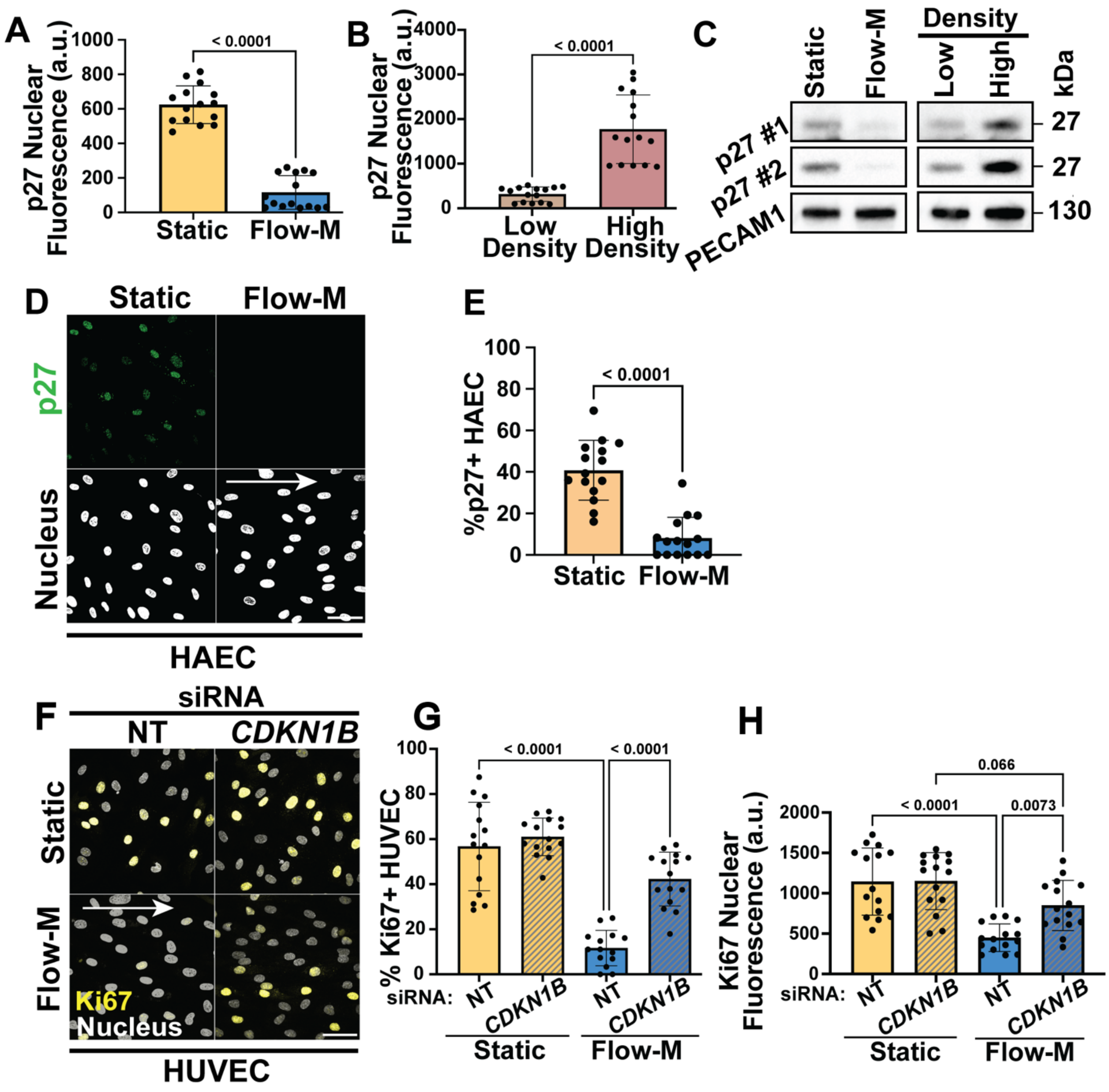


**Supplemental Figure 3. Cell cycle inhibitor p27 expression differs with quiescence stimulus.**

**(A)** Quantification of p27 nuclear fluorescence intensity in indicated conditions. n=3 replicates, 5 images averaged per condition per replicate. **(B)** Quantification of p27 nuclear fluorescence intensity in indicated conditions. n=3 replicates, 5 images averaged per condition per replicate. **(C)** Western blot of p27 expression under indicated conditions, and with indicated antibodies. p27 Ab #1 (Cell Signaling) and p27 Ab #2 (Santa Cruz). **(D)** Representative images of HAEC under indicated conditions stained for p27 (green) and DAPI (white, nuclear mask). Scale bar, 50 µm. White arrow, flow direction. **(E)** Quantification of HAEC p27+ cells under indicated conditions, n=3 replicates, 5 images per condition per replicate. **(F)** Representative images of HUVEC with indicated siRNA and treatments. Endothelial cells stained with Ki67 (yellow, proliferation marker) and DAPI (gray, nucleus mask). Scale bar 50 µm. White arrow, flow direction. **(G)** Quantification of percent Ki67+ cells. n=3 replicates, 5 images per condition per replicate. **(H)** Quantification of nuclear fluorescence intensity of Ki67. n=3 replicates, 5 averaged images per condition per replicate. Statistics, student's two-tailed *t*-test (A-B, E) and one-way ANOVA with Tukey's multiple comparisons test (G-H).

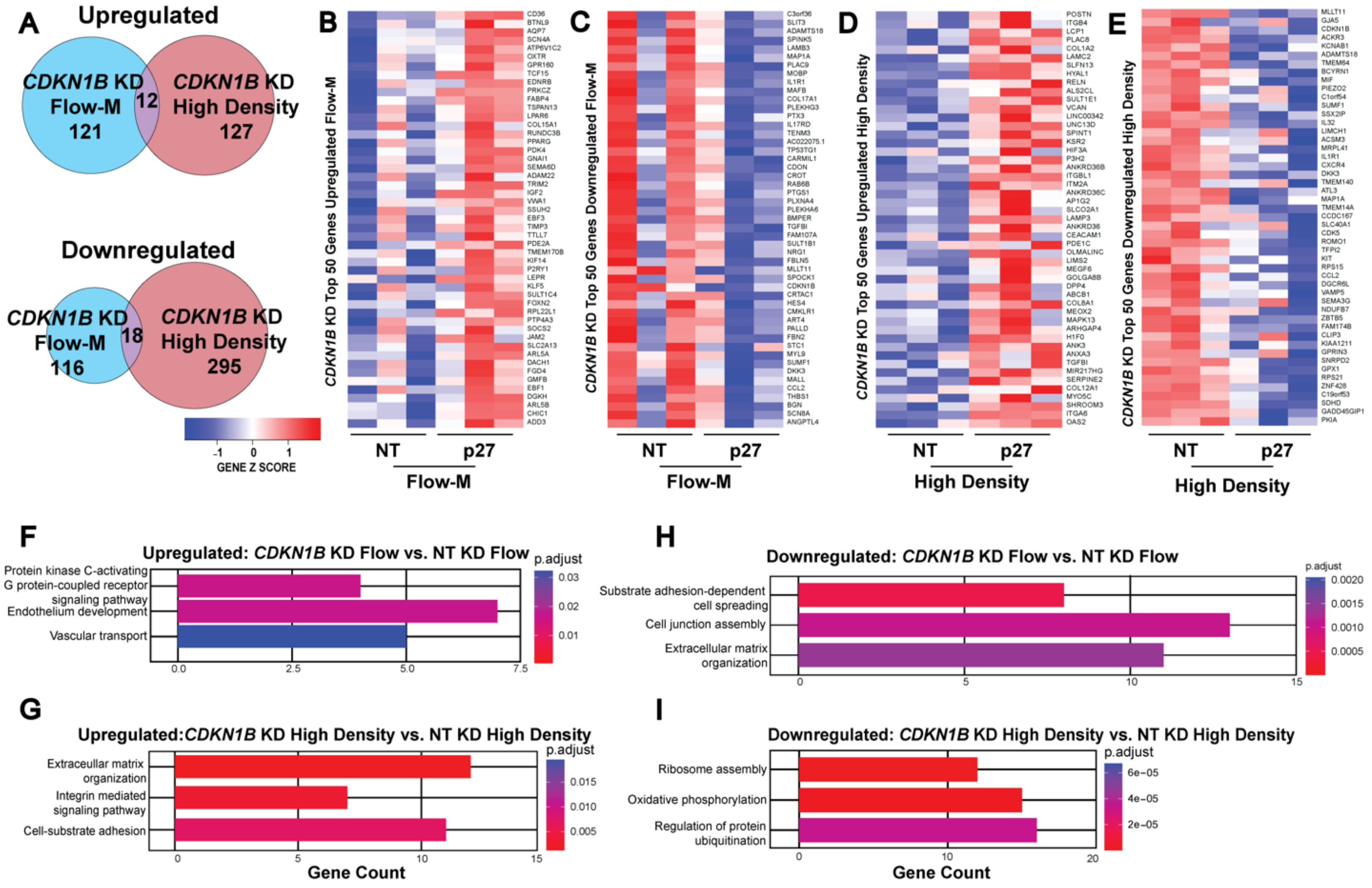


**Supplemental Figure 4. p27 depletion leads to distinct transcriptional changes dependent on quiescence stimulus.**

**(A)** Venn diagrams showing overlap of HUVEC genes differentially regulated in indicated conditions and for *CDKN1B* KD compared to NT. **(B-E)** Heatmaps showing relative expression levels of genes differentially regulated (top 50 by fold change and p-value) in response to indicated conditions. **(F-I)** GO analysis performed on differentially expressed genes from bulk RNA-seq data comparing indicated conditions/treatments. Representative biological processes GO terms significantly enriched (P adjusted < 0.1) in differentially regulated genes are shown.

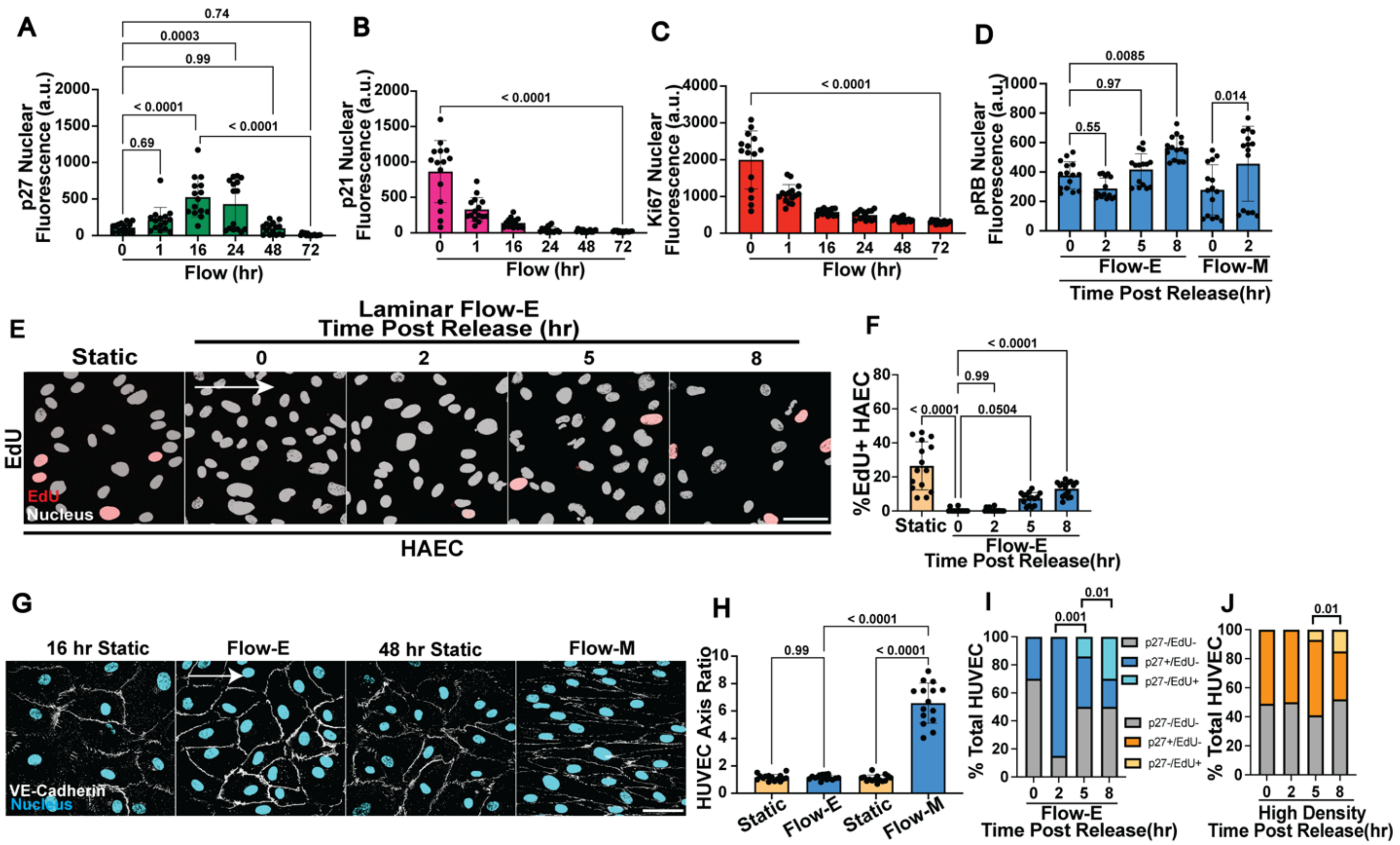


**Supplemental Figure 5. Cell cycle inhibitor and Ki67 expression intensity varies with flow and correlates with flow-mediated endothelial cell alignment.**

**(A)** Quantification of p27 nuclear fluorescence intensity under indicated conditions. n=3 replicates, 5 averaged images per condition per replicate. **(B)** Quantification of p21 nuclear fluorescence intensity under indicated conditions. n=3 replicates, 5 averaged images per condition per replicate. **(C)** Quantification of Ki67 nuclear fluorescence intensity under indicated conditions. n=3 replicates, 5 averaged images per condition per replicate. **(D)** Quantification of pRB nuclear fluorescence intensity under indicated conditions. n=3 replicates, 5 averaged images per condition per replicate. **(E)** Representative images of HAEC under static (non-flow) or Flow-E conditions with EdU incorporation and fixation at indicated times post Flow-E release. Cells stained for DAPI (white, nuclear mask) and EdU (red, S-phase), Scale bar, 50 µm. White arrow, flow direction. **(F)** Quantification of EdU+ cells under indicated conditions. n=3 replicates, 5 images per condition per replicate. **(G)** Representative images of HUVEC stained with VE-cadherin (white, junction marker) and DAPI (blue, nucleus) under indicated conditions. Scale bar, 50 µm. White arrow, flow direction. **(H)** Cell axis ratio quantification under indicated conditions. n=3 replicates, 5 images per condition per replicate. **(I)** % HUVEC under Flow-E release with p27-/EdU-, p27+/EdU-, and p27-/EdU+ incorporation. **(J)** % HUVEC under high density release with p27-/EdU-, p27+/EdU-, and p27-/EdU+ incorporation. Statistics, one-way ANOVA with Tukey's multiple comparisons test (A-D, F, H) and $C^2$ test (I-J).

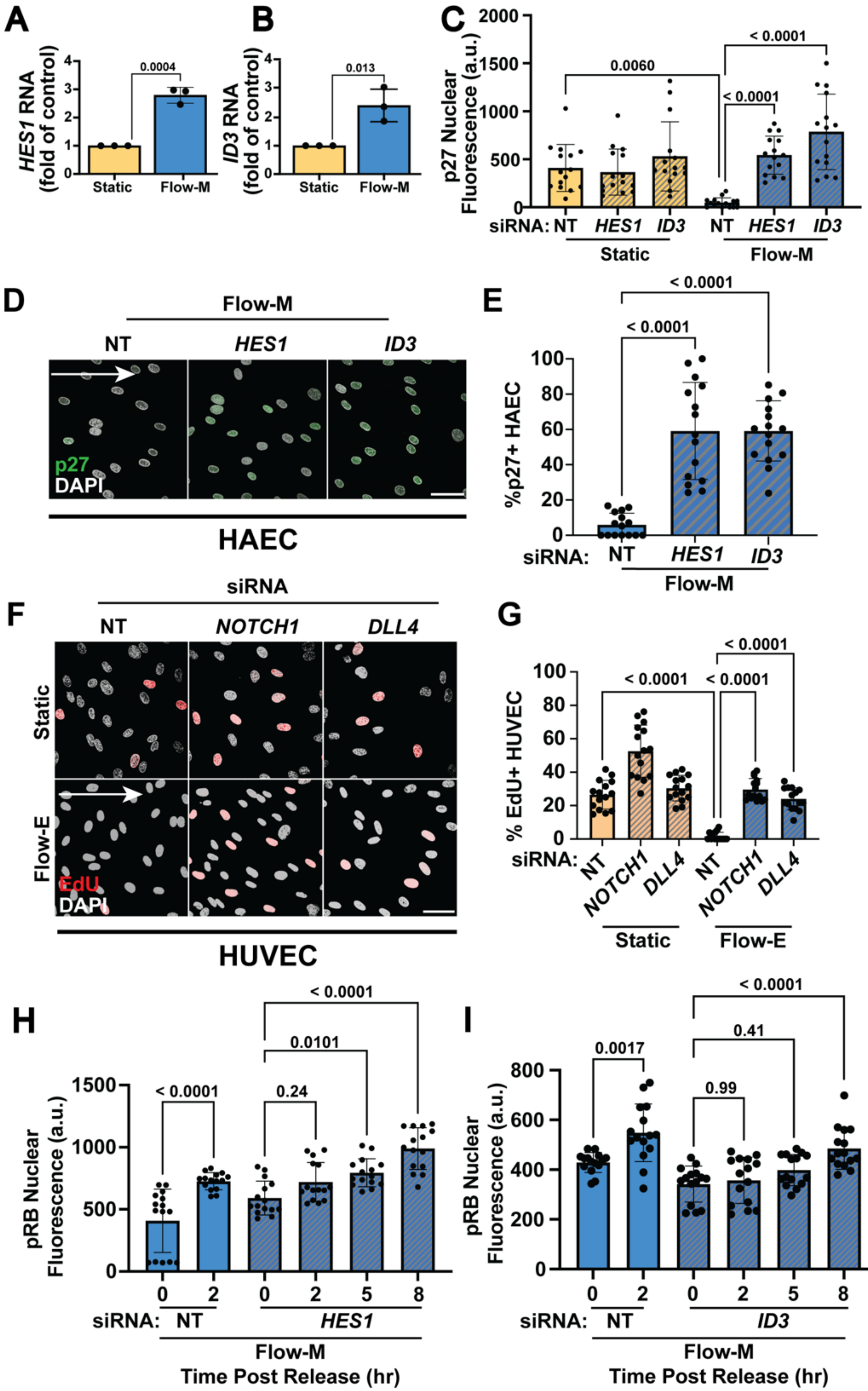
A
HES1 RNA (fold of control)
0.0004
Static
Flow-M
B
ID3 RNA (fold of control)
0.013
Static
Flow-M
C
p27 Nuclear Fluorescence (a.u.)
0.0060
< 0.0001
< 0.0001
siRNA: NT HES1 ID3 NT HES1 ID3
Static
Flow-M
D
Flow-M
NT
HES1
ID3
p27
DAPI
HAEC
E
%p27+ HAEC
< 0.0001
< 0.0001
siRNA: NT HES1 ID3
Flow-M
F
siRNA
NT
NOTCH1
DLL4
Static
Flow-E
EdU
DAPI
HUVEC
G
% EdU+ HUVEC
< 0.0001
< 0.0001
< 0.0001
siRNA: NT NOTCH1 DLL4 NT NOTCH1 DLL4
Static
Flow-E
H
pRB Nuclear Fluorescence (a.u.)
< 0.0001
0.24
0.0101
< 0.0001
0 2 0 2 5 8
siRNA: NT HES1
Flow-M
Time Post Release (hr)
I
pRB Nuclear Fluorescence (a.u.)
0.0017
0.99
0.41
< 0.0001
0 2 0 2 5 8
siRNA: NT ID3
Flow-M
Time Post Release (hr)

**Supplemental Figure 6. Nuclear fluorescence intensity of markers and changes with *HES1* or *ID3* depletion.**

**(A)** RT-qPCR for *HES1* expression under indicated conditions. n=3 replicates. **(B)** RT-qPCR for *ID3* expression under indicated conditions, n=3 replicates. **(C)** Quantification of p27 nuclear fluorescence intensity with indicated siRNA treatments and conditions. n=3 replicates, 5 averaged images per condition per replicate. **D)** Representative images of HAEC under indicated conditions and with indicated siRNA treatment. Cells stained for p27 (green) and DAPI (white, nuclear mask). Scale bar, 50 µm. White arrow, flow direction. **(E)** Quantification of p27+ cells under indicated conditions and treatments. n=3 replicates, 5 images per condition per replicate. **(F)** Representative images of HUVEC under indicated conditions and siRNA treatments. Cells were labeled with EdU (red, S-phase) and stained for DAPI (white, nuclear mask). Scale bar, 50 µm. White arrow, flow direction. **(G)** Quantification of EdU+ cells with indicated conditions. n=3 replicates, 5 images per condition per replicate. **(H)** Quantification of pRB nuclear fluorescence intensity under indicated siRNA treatments and conditions. n=3 replicates, 5 averaged images per condition per replicate. **(I)** Quantification of pRB nuclear fluorescence intensity under indicated siRNA treatments and conditions. n=3 replicates, 5 averaged images per condition per replicate. Statistics, student's two tailed t-test (A-B) and one-way ANOVA with Tukey's multiple comparisons test (C, E, G-I).

**Supplementary Table S1. Genes Used to Generate Endothelial Quiescence Score**

| Gene | Expression | Gene | Expression | Gene | Expression |
|---|---|---|---|---|---|
| MCM5 | downregulated | CASP8AP2 | downregulated | HN1 | downregulated |
| PCNA | downregulated | USP1 | downregulated | CDC20 | downregulated |
| TYMS | downregulated | CLSPN | downregulated | TTK | downregulated |
| FEN1 | downregulated | POLA1 | downregulated | CDC25C | downregulated |
| MCM2 | downregulated | CHAF1B | downregulated | KIF2C | downregulated |
| MCM4 | downregulated | BRIP1 | downregulated | RANGAP1 | downregulated |
| RRM1 | downregulated | E2F8 | downregulated | NCAPD2 | downregulated |
| UNG | downregulated | HMGB2 | downregulated | DLGAP5 | downregulated |
| GINS2 | downregulated | CDK1 | downregulated | CDCA2 | downregulated |
| MCM6 | downregulated | NUSAP1 | downregulated | CDCA8 | downregulated |
| CDCA7 | downregulated | UBE2C | downregulated | ECT2 | downregulated |
| DTL | downregulated | BIRC5 | downregulated | KIF23 | downregulated |
| PRIM1 | downregulated | TPX2 | downregulated | HMMR | downregulated |
| UHRF1 | downregulated | TOP2A | downregulated | AURKA | downregulated |
| MLF1IP | downregulated | NDC80 | downregulated | PSRC1 | downregulated |
| HELLS | downregulated | CKS2 | downregulated | ANLN | downregulated |
| RFC2 | downregulated | NUF2 | downregulated | LBR | downregulated |
| RPA2 | downregulated | CKS1B | downregulated | CKAP5 | downregulated |
| NASP | downregulated | MKI67 | downregulated | CENPE | downregulated |
| RAD51AP1 | downregulated | TMPO | downregulated | CTCF | downregulated |
| GMNN | downregulated | CENPF | downregulated | NEK2 | downregulated |
| WDR76 | downregulated | TACC3 | downregulated | G2E3 | downregulated |
| SLBP | downregulated | FAM64A | downregulated | GAS2L3 | downregulated |
| CCNE2 | downregulated | SMC4 | downregulated | CBX5 | downregulated |
| UBR7 | downregulated | CCNB2 | downregulated | CENPA | downregulated |
| POLD3 | downregulated | CKAP2L | downregulated | GABPB2 | downregulated |
| MSH2 | downregulated | CKAP2 | downregulated | MAD2L1 | downregulated |
| ATAD2 | downregulated | AURKB | downregulated | VEGFC | downregulated |
| RAD51 | downregulated | BUB1 | downregulated | RB1 | upregulated |
| RRM2 | downregulated | KIF11 | downregulated | PEPD | upregulated |
| CDC45 | downregulated | ANP32E | downregulated | CDKN1A | upregulated |
| CDC6 | downregulated | TUBB4B | downregulated | CDKN1C | upregulated |
| EXO1 | downregulated | GTSE1 | downregulated | KMT5A | upregulated |
| TIPIN | downregulated | KIF20B | downregulated | MAPK14 | upregulated |
| DSCC1 | downregulated | HJURP | downregulated | ZNF124 | upregulated |
| BLM | downregulated | CDCA3 | downregulated | HES1 | upregulated |

S

**Supplementary Table S2. Bulk RNA Sequencing Mapping Rate**

| Sample | Total Counts | Mapped Counts | Mapped Counts (%) |
|---|---|---|---|
| NT_STAT1 | 22755743 | 20783635 | 91.3 |
| NT_STAT2 | 19339936 | 17604990 | 91.0 |
| NT_STAT3 | 19787509 | 18176842 | 91.9 |
| NT_FLOW1 | 21604345 | 19966597 | 92.4 |
| NT_FLOW2 | 22552162 | 20448912 | 90.7 |
| NT_FLOW3 | 21098509 | 19220279 | 91.1 |
| p27_STAT1 | 19933252 | 17889154 | 89.7 |
| p27_STAT2 | 14798902 | 13372009 | 90.4 |
| p27_STAT3 | 18964619 | 17181169 | 90.6 |
| p27_FLOW1 | 20814162 | 18963329 | 91.1 |
| p27_FLOW2 | 27363377 | 25100897 | 91.7 |
| p27_FLOW3 | 31026010 | 27242573 | 87.8 |
| NT_LD1 | 21548406 | 19470718 | 90.4 |
| NT_LD2 | 18915090 | 16447249 | 87.0 |
| NT_LD3 | 23546196 | 21639660 | 91.9 |
| NT_HD1 | 19596251 | 18071656 | 92.2 |
| NT_HD2 | 19383071 | 17828106 | 92.0 |
| NT_HD3 | 26543935 | 24379725 | 91.8 |
| p27_LD1 | 26198842 | 24291108 | 92.7 |
| p27_LD2 | 20686749 | 18790131 | 90.8 |
| p27_LD3 | 21395446 | 19699259 | 92.1 |
| p27_HD1 | 20567925 | 19008271 | 92.4 |
| p27_HD2 | 19622332 | 17806879 | 90.7 |
| p27_HD3 | 17857053 | 16355030 | 91.6 |

### Major Resources Table

#### Animals (in vivo studies)

| Species | Vendor or Source | Background Strain | Sex | Persistent ID / URL |
|---|---|---|---|---|
| *Danio Rerio* | Wiebke Herzog Lab | AB | N/A | N/A |

#### Genetically Modified Animals

| | Species | Vendor or Source | Background Strain | Other Information | Persistent ID / URL |
|---|---|---|---|---|---|
| **Parent - Male** | *Danio Rerio* | Victoria Bautch Lab | AB | *cdkn1bb* deletion | https://zfin.org/ZDB-ALT-231024-2 |
| **Parent - Female** | *Danio Rerio* | Victoria Bautch Lab | AB | *ckdn1bb* deletion | https://zfin.org/ZDB-ALT-231024-2 |

#### Antibodies

| Target antigen | Vendor or Source | Catalog # | Working Concentration | Persistent ID / URL |
|---|---|---|---|---|
| p27 | Cell Signaling | 3686S | IF, WB: 1:1000 | https://www.cellsignal.com/products/primary-antibodies/p27-kip1-d69c12-xp-rabbit-mab/3686 |
| p27 | Santa Cruz | sc-1641 | WB: 1:1000 | https://www.scbt.com/p/p27-antibody-f-8 |
| p21 | Santa Cruz | sc-6246 | IF: 1:50 | https://www.scbt.com/p/p21-antibody-f-5 |
| Ki67 | Abcam | ab15580 | IF: 1:1000 | https://www.abcam.com/products/primary-antibodies/ki67-antibody-ab15580.html |
| Ki67 | ThermoFisher | 14-5698-82 | IF: 1:1000 | https://www.thermofisher.com/antibody/product/Ki-67-Antibody-clone-SolA15-Monoclonal/14-5698-82 |
| VE-Cadherin | Santa Cruz | sc-9989 | IF: 1:100 | https://www.scbt.com/p/ve-cadherin-antibody-f-8?gad_source=1&gclid=Cj0KCQiA7OqrBhD9ARIsAK3UXh2GA0sNEM06ICNotaVduI2oO3jDlSCbm_N_iuY_80rxTEYLKIMovOwaAiT7EALw_wcB |
| PECAM1 | Cell Signaling | 3528S | WB: 1:1000 | https://www.cellsignal.com/products/primary-antibodies/cd31-pecam-1-89c2-mouse-mab/3528 |
| Goat anti rabbit IgG (H+L) Secondary Antibody, Alexa Fluor 488 | ThermoFisher | A11034 | IF: 1:1000 | https://www.thermofisher.com/antibody/product/Goat-anti-Rabbit-IgG-H-L-Highly-Cross-Adsorbed-Secondary-Antibody-Polyclonal/A-11034 |
| Goat anti rabbit IgG (H+L) Secondary Antibody, Alexa Fluor 594 | ThermoFisher | A11012 | IF: 1:1000 | https://www.thermofisher.com/antibody/product/Goat-anti-Rabbit-IgG-H-L-Cross-Adsorbed-Secondary-Antibody-Polyclonal/A-11012 |
| Goat anti rabbit IgG (H+L) Secondary Antibody, Alexa Fluor 647 | ThermoFisher | A21245 | IF: 1:1000 | https://www.thermofisher.com/antibody/product/Goat-anti-Rabbit-IgG-H-L-Highly-Cross-Adsorbed-Secondary-Antibody-Polyclonal/A-21245 |
| Donkey anti-Rabbit IgG (H+L) Highly Cross-Adsorbed Secondary Antibody, HRP | ThermoFisher | A16035 | WB: 1:10000 | https://www.thermofisher.com/antibody/product/Donkey-anti-Rabbit-IgG-H-L-Highly-Cross-Adsorbed-Secondary-Antibody-Polyclonal/A16035 |

| | | | | |
|---|---|---|---|---|
| Donkey anti-Mouse IgG (H+L) Highly Cross-Adsorbed Secondary Antibody, HRP | ThermoFisher | A16011 | WB: 1:10000 | https://www.thermofisher.com/antibody/product/Donkey-anti-Mouse-IgG-H-L-Secondary-Antibody-Polyclonal/A16011 |
| Phospho-Rb (Ser807/811) (D20B12) XP® Rabbit mAbAlexa Fluor 647 Conjugate | Cell Signaling | 8974S | IF: 1:1000 | https://www.cellsignal.com/products/antibody-conjugates/phospho-rb-ser807-811-d20b12-xp-rabbit-mab-alexa-fluor-647-conjugate/8974 |
| DAPI | Sigma | 10236276001 | IF: 1:1000 | https://www.sigmaaldrich.com/US/en/product/roche/10236276001 |

**Cultured Cells**

| Name | Vendor or Source | Sex (F, M, or unknown) | Persistent ID / URL |
|---|---|---|---|
| HUVEC | Lonza | Pooled Donor | https://bioscience.lonza.com/lonza_bs/US/en/Primary-and-Stem-Cells/p/000000000000184665/HUVEC-%E2%80%93-Human-Umbilical-Vein-Endothelial-Cells%2C-Pooled%2C-in-EGM%E2%84%A2-2 |
| HAEC | Lonza | Pooled Doner | https://bioscience.lonza.com/lonza_bs/US/en/Primary-and-Stem-Cells/p/000000000000184983/HAEC-%E2%80%93-Human-Aortic-Endothelial-Cells |

**Data & Code Availability**

| Description | Source / Repository | Persistent ID / URL |
|---|---|---|
| Bulk RNAseq Flow and Contact Inhibition *in vitro* | GSE213323 | Available upon request |
| scRNAseq mouse neonatal ear skin | GSE216594 | Available upon request |

**siRNAs**

| siRNA | Company | Catalog # | Target Sequence (5'-> 3') |
|---|---|---|---|
| *CDKN1B* (p27) | Dharmacon | L-003472-00-0005 | CAAACGUGCGAGUGUCUAA<br>GCAGCUUGCCCGAGUUCUA<br>ACGUAAACAGCUCGAAUUA<br>GCAAUGCGCAGGAAUAAGG |
| *CDKN1B* (p27) | Dharmacon | J-003472-05-0005 | CAAACGUGCGAGUGUCUAA |
| *ID3* | Dharmacon | L-009905-00-0005 | GCACUCAGCUUAGCCAGGU<br>GAACGCAGUCUGGCCAUCG<br>GGGAACUGGUACCCGGAGU<br>GGAAGGUGACUUUCUGUAA |
| *ID3* | Dharmacon | J-009905-05-0005 | GCACUCAGCUUAGCCAGGU |
| *HES1* | Dharmacon | L-007770-02-0005 | ACGAAGAGCAAGAAUAAAU<br>AGGCUGGAGAGGCGGCUAA<br>UCAACACGACACCGGAUAA<br>ACUGCAUGACCCAGAUCAA |
| *HES1* | Dharmacon | J-007770-21-0005 | ACGAAGAGCAAGAAUAAAU |
| *NOTCH1* | Santa Cruz | sc-36095 | N/A |
| *DLL4* | Life Tech | s29214 | GGUACCUUCUCGCUCAUCAtt |
| *Non-Targeting* 1 | Dharmacon | D-001810-10-20 | N/A |
| *Non-Targeting* 2 | ThermoFisher | 4390844 | N/A |
| *Non-Targeting* 3 | ThermoFisher | 4390847 | N/A |

**qPCR Primers**

| Gene | Species | Primer Sequence (5'-> 3') |
|---|---|---|
| *CDKN1B* Forward | Human | ATCACAAACCCCTAGAGGGCA |
| *CDKN1B* Reverse | Human | GGGTCTGTAGTAGAACTCGGG |
| *ID3* Forward | Human | CAGCTTAGCCAGGTGGAAATCC |

| | | |
|---|---|---|
| *ID3* Reverse | Human | GTCGTTGGAGATGACAAGTTCCG |
| *HES1* Forward | Human | GGAAATGACAGTGAAGCACCTCC |
| *HES1* Reverse | Human | GAAGCGGGTCACCTCGTTCATG |
| *b Actin* Forward | Human | GATGGCCACGGCTGCTTC |
| *b Actin* Reverse | Human | TGCCTCAGGGCAGCGGAA |
| *cdkn1bb* Forward | Zebrafish | CCGTCGGCATGTCGAAACCT |
| *cdkn1bb* Reverse | Zebrafish | GCGTGTGCGTGGAAAAGTCG |
| *mki67* Forward | Zebrafish | GAACACCCTCGACAACGCCA |
| *mki67* Reverse | Zebrafish | GACCCACCACTTCGCCTTCC |
| *ccdn1* Forward | Zebrafish | GGCGTCATGGGACAAGAGGG |
| *ccnd1* Reverse | Zebrafish | CCTGGCCAGCGATAACTCGG |
| *pcna* Forward | Zebrafish | GACACGCTGGCACTGGTCTT |
| *pcna* Reverse | Zebrafish | TGCAGATGCGGGCAAACTCA |
| *gapdh* Forward | Zebrafish | TGACCTGATGGCACACATGG |
| *gapdh* Reverse | Zebrafish | TGGGAGAATGGTCGCGTATC |
| *b2m* Forward | Zebrafish | GCCTTCACCCCAGAGAAAGG |
| *b2m* Reverse | Zebrafish | GCGGTTGGGATTTACATGTTG |
